# Dynamical processing of orientation precision in the primary visual cortex

**DOI:** 10.1101/2021.03.30.437692

**Authors:** Hugo J. Ladret, Nelson Cortes, Lamyae Ikan, Frédéric Chavane, Christian Casanova, Laurent U. Perrinet

## Abstract

In our daily visual environment, the primary visual cortex (V1) processes distributions of oriented features as the basis of our visual computations. Changes of the global, median orientation of such inputs form the basis of our canonical knowledge about V1. However, another overlooked but defining characteristic of these sensory variables is their precision, which characterizes the level of variance in the input to V1. Such variability is an intrinsic part of natural images, yet it remains unclear if and how V1 accounts for the changes in orientation precision to achieve its robust orientation recognition performances. Here, we used naturalistic stimuli to characterize the response of V1 neurons to quantified variations of orientation precision. We found that about thirty percent of the recorded neurons showed a form of invariant responses to input precision. While feedforward mechanisms failed to account for the existence of these resilient neurons, neuronal competition within V1 explained the extent to which a neuron is invariant to precision. Using a decoding algorithm, we showed that the existence of such neurons in the population response of V1 can serve to encode both the orientation and its precision in the V1 population activity, which improves the robustness of the overall neural code. These precision-specific neurons operate with slow recurrent cortical dynamics, which supports the notion of predictive precision-weighted processes in V1.

## Introduction

Selectivity to the orientation of stimuli is an archetypal feature of neurons in the mammalian primary visual cortex (V1) (34). In more than 60 years of investigations, stimuli such as oriented gratings have uncovered many filter-like properties in V1 (61). Fundamentally, these low-complexity stimuli are limited in their capability to explain how the rich cortical dynamics involved in the vision of natural images (20, 82) are driven by the complex content of our everyday visual scenery (72). Rather than gratings or Gabor filters, natural inputs to the visual system are distributions of orientations (Supplementary Figure 1). Yet, most of our current understanding of orientation processing still comes from studies using single-orientation stimuli, i.e. made of an exaggeratedly precise orientation distribution. The effect of orientation precision on orientation selectivity is therefore a major gap that needs to be addressed.

Furthermore, the precision of a sensory variable is of central importance to empirical Bayesian inference, in which it weights the relative influence of the likelihood with respect to a prior when computing the estimation of the posterior distribution. In predictive coding (PC) terms (21, 22), precision allows the brain to balance between its own internal representations (predictions) and its new sensory inputs (prediction errors) to accurately predict the states in the environment. Practically, given a low-precision image of a lion skulking through the savanna, a predictive coding system that seeks to avoid this predator should (relatively to a high-precision image) rely more on its “lion” internal prediction than on its “savanna grass” sensory input to produce an optimal behavior. Given the mounting evidence that PC captures the behavior of the visual system (7, 63, 69), it is physiologically likely and mathematically required by PC - that the first level of the visual hierarchy, V1, possesses a mechanism which can account for the precision of sensory inputs.

Previous investigations have shown that orientation precision causes non-linear tuning modulations of single V1/V2 neurons (26), which is accounted for by models of recurrent cortical activity of V1 (2, 3, 42) and correlates with psychophysical performances (30, 31, 60). These converging observations have led to the notion that population variability can be used to encode sensory feature precision (32), yet the detailed processes that would be involved in such a population code remain by and large nonestablished. Here, we used natural-like stimuli to study the encoding of the precision of oriented inputs in V1. We quantified the precision-response relationships of single neurons, which shed light on two types of precision-modulated activity profiles. Some neurons exhibited “precision-resilient” activity, which, as opposed to “precision-vulnerable” neurons, remained well-tuned even at minimal input precision. These resilient neurons reached a peak spiking activity late after stimulation onset, and had an optimal tuning delay which depended on the input’s precision. A phenomenological recurrent neural network showed that varying the strength of lateral cortical inhibition can explain the existence of these two categories of neurons.

This compelling evidence for the involvement of recurrent connectivity in precision responses led us to investigate how precision modulates the overall population activity in V1. Using a biologically plausible neuronal decoder (4), we found that orientation could be robustly inferred from the population activity at each level of precision. We found that precision and orientation are co-encoded only by resilient neurons, significantly improving the orientation encoding of the neural population. This encoding involves specific neural dynamics, in which the cortical processing time and the population variability are directly related to the input precision, resulting in temporally segregated neural representations of precision within the same neural network. Taken together, these results suggest that the precision of sensory inputs is actively processed in V1. Additionally, slow non-canonical dynamics and neuronal subpopulations dedicated to the processing of input precision in V1 are compelling experimental traces of a predictive process occurring in our recordings.

## Results

### Single neuron modulations by orientation precision

Single-unit activity of 249 V1 neurons was recorded in three anesthetized cats. Orientation selectivity was measured using band-pass filtered white noise images called Motion Clouds (45), whose underlying generative framework allowed to finely control the orientation distribution of each visual stimulus. We focused on varying two parameters of the Motion Clouds: the median orientation of the distribution, *θ*, and the bandwidth of the distribution *B*_*θ*_ (Supplementary Figure 1b). This parameter is a measure of the standard deviation *σ* of the distribution, which is directly related to the precision 1*/σ*^2^. Hence, increasing *B*_*θ*_ corresponds to decreasing the precision of the oriented input to V1. An additional benefit from using Motion Clouds is their stationarity in the spatial domain, which removes any potential secondorder correlation found in natural images, further allowing an isolation of the effect of precision on orientation selectivity (Supplementary Figure 1b). Moreover, Motion Clouds conform to the statistics of natural images, namely the 1*/f* ^2^ distribution of the power spectrum (19).

We generated Motion Clouds at 12 evenly spaced values of *θ* (steps of 15°) and 8 evenly distributed values of *B*_*θ*_ (starting at ≈ 0°, ending at 36°) and measured the orientation selectivity of single V1 neurons. As precision decreased (i.e. *B*_*θ*_ increased), almost all tuning curves remained centered on the same preferred orientation (98.8% units, *p* < 0.05, Wilcoxon signed-rank test) and diminished in peak amplitude (95.1% units, *p* < 0.05, Wilcoxon signedrank test, 73.1% mean amplitude decrease). Only 28.5% of the recorded neurons were still tuned when orientation precision reached *B*_*θ*_ = 36.0° (*p* < 0.05, Wilcoxon signed-rank test). Hence, a reduction in the precision of oriented inputs to V1 caused a decrease in single neuron tuning. This precision modulation unfolded heterogeneously across neurons, as illustrated by two example tuning curves (Figure 1b). Neuron Ma006 was no longer tuned to stimuli of *B*_*θ*_ = 36.0° (*W* = 171.0, *p* = 0.24, Wilcoxon signed-rank test, firing rate of preferred vs orthogonal orientation), contrary to neuron Tv001 which remained orientation selective even at the lowest input precision used here (*W* = 22.5, *p* = 10^−6^).

**Figure 1.**
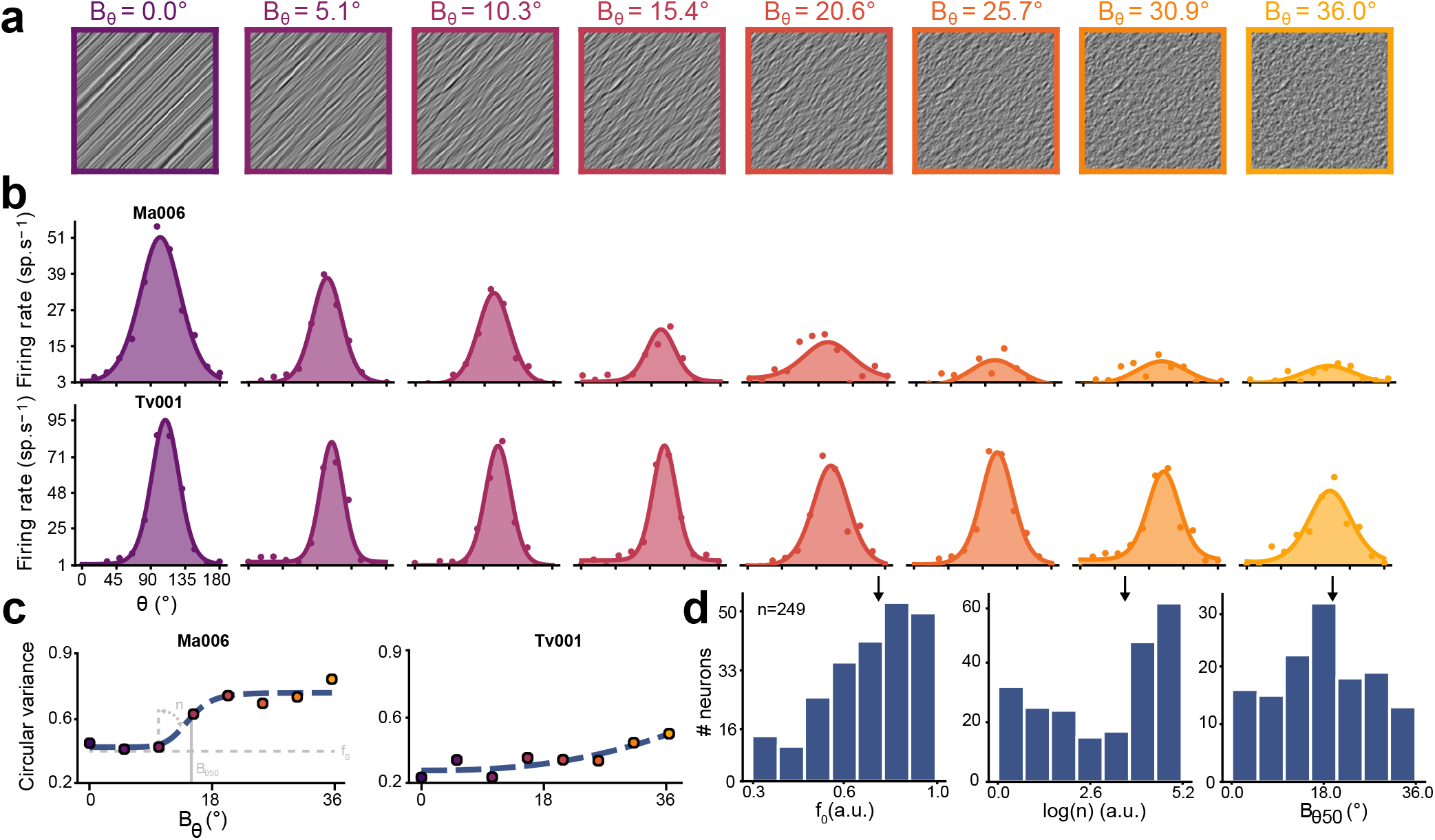
Single neuron responses to variations of orientation precision. Additional examples are provided in Supplementary Figure 2. (**a**) Examples of Motion Clouds, oriented at 45° relative to the vertical axis. Neural activity was elicited with Motion Clouds of 12 different orientations *θ* and 8 orientation precision *B*_*θ*_, drifting orthogonally to their main orientation. (**b**) Tuning curves of two examples neurons (labelled ‘Ma006’ and ‘Tv001’) stimulated with Motion Clouds of increasing *B*_*θ*_ from left to right, i.e. of decreasing orientation precision. Colored dots represent the mean firing rate across trials (baseline subtracted, 300 ms average), with lines indicating a fitted von Mises function. (**c**) Precision-response functions (PRFs), assessing the changes of orientation tuning as measured by the Circular Variance (CV, colored dots) as a function of *B*_*θ*_, fitted with Naka-Rushton (NKR) functions (dashed curves). The effect of NKR parameters are illustrated (in gray) for neuron Ma006’s PRF. The parameters for Ma006’s PRF are *f*_0_ = 0.4, *n* = 8.4, *B*_*θ*50_ = 14.7°. For Tv001, they are *f*_0_ = 0.3, *n* = 2.4, *B*_*θ*50_ = 36.0°. (**d**) Distribution of the NKR parameters for all recorded neurons. Median values of the distributions are indicated by a black arrow (*f*_0_ = 0.75, log(*n*) = 3.6, *B*_*θ*50_ = 19.2).

These two distinctive neuronal behaviors were further evidenced by measuring the variation of the goodness of tuning (circular variance, CV) as a function of orientation precision *B*_*θ*_. Such “precision-response functions” (PRFs) uncovered various degrees of non-linearity in the population (Figure 1c) and were readily captured by a Naka-Rushton function (53) (Supplementary Figure 3). This function characterized the response to variations of precision with three parameters: *f*_0_, the goodness of orientation tuning for most precise inputs ; *n*, the degree of non-linearity of the PRF and *B*_*θ*50_, the level of input precision at which a neuron’s response transitioned from tuned to untuned state. Overall, the PRF parameters were highly heterogeneous through all the recorded neurons (Figure 1d). To better understand the principles underlying this response heterogeneity, we separated the recorded units in two groups. Neurons which remained significantly orientation tuned at lowest precision (*B*_*θ*_ = 36.0°) were labeled “resilient neurons” (71 neurons, 28.51 % of recorded units), while the rest of the population, which was no longer tuned at lowest precision, were labeled “vulnerable neurons”. This is an artificial split that serves to illustrate how the non-linearity of precision responses can exist, by separating the most extreme behaviors (those of resilient neurons) from the rest of the recordings. These two groups exhibited different dynamics: the maximum firing rate of resilient neurons was delayed compared to vulnerable ones, regardless of the input’s precision (Figure 2g,a, *U* = 8256.0, *p* = 8.10^−5^ for *B*_*θ*_ = 0.0° and *U* = 7378.0, *p* = 0.01 for *B*_*θ*_ = 36.0°, Mann-Whitney U test). Resilient neurons displayed a delayed time to maximum orientation tuning which changed with input precision (Figure 2h,b, *U* = 1815.0, *p* = 0.002, Mann-Whitney U test). Both groups of neurons also differed in their PRFs parameters. While their circular variances were overlapping (Figure 2c,d), resilient neurons showed significantly lower *f*_0_ (Figure 2e, *U* = 4663.0, *p* = 0.004, MannWhitney U test) and *n* (Figure 2f, *U* = 5352.5, *p* = 0.04, Mann-Whitney U test). Respectively, this corresponds to a better tuning to precise oriented inputs and reduced variations of tuning when precision is decreasing, compared to vulnerable neurons. Altogether, resilient neurons exhibited sharper orientation tuning and slower dynamics, which are distinctive features of supragranular neurons (65, 66). This correlates with the location of the recorded units, which predominantly places resilient neurons in supragranular layers (Figure 2i). In terms of precision responses, resilient neurons’ more linear PRFs gives them increased dynamic range, resulting in a better orientation coding in across a wide range of precision (43, 44). It should be noted that these two groups of neurons showed no difference in their motion direction selectivity index (*U* = 6569.5, *p* = 0.62, Mann-Whitney U test). Hence, the presence of these resilient versus vulnerable neurons is not explained by motion integration, which here had constant precision (drift coherence), as opposed to orientation precision.

**Figure 2.**
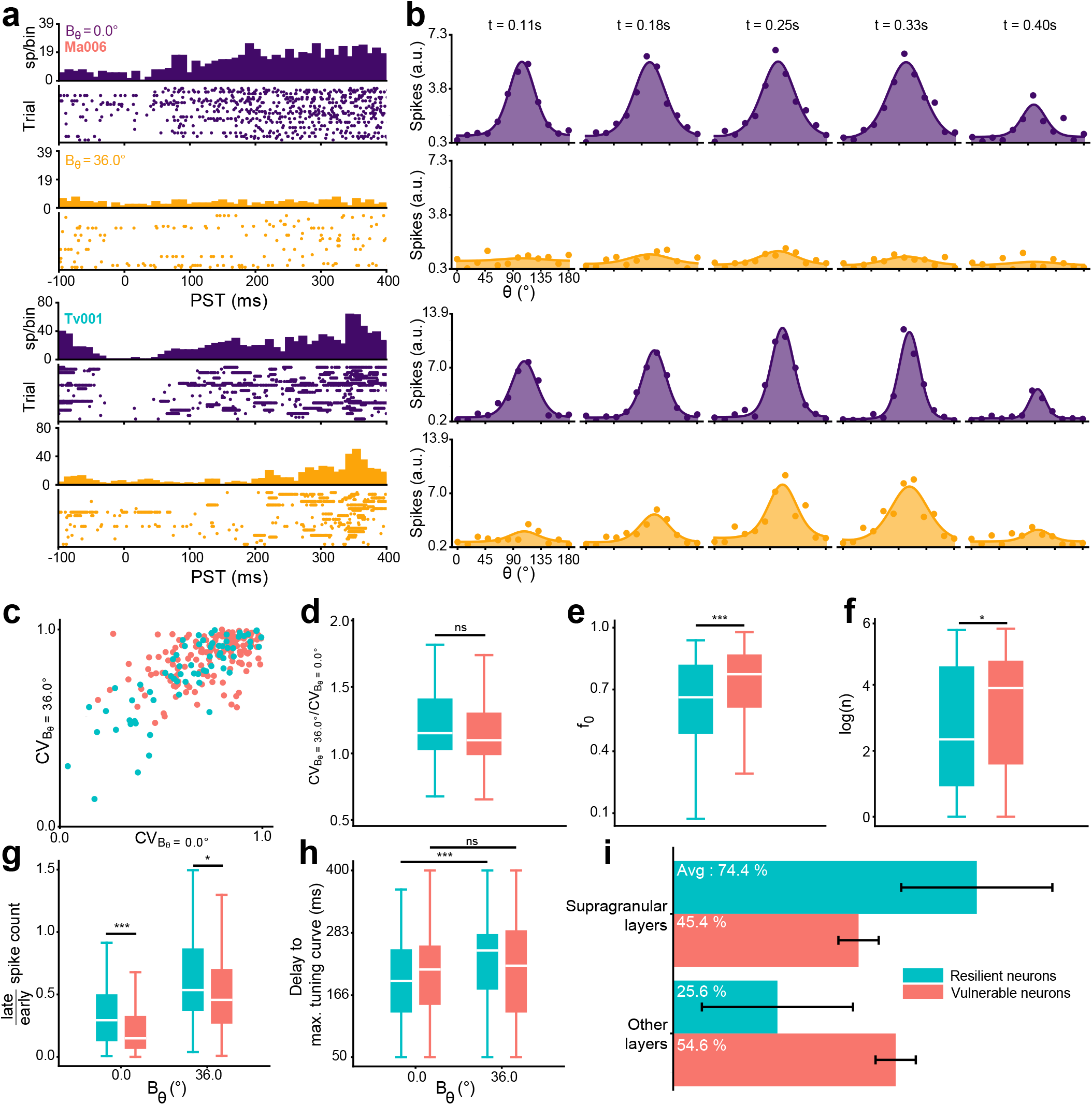
Dynamical properties of the neuronal responses. Neuron labels are colored in orange for neurons untuned to stimuli with *B*_*θ*_ = 36.0° (“vulnerable neurons”), and conversely colored in cyan when still significantly tuned to the same lowest-precision stimuli (“resilient neurons”). Additional examples are provided in Supplementary Figure 4. (**a**) Peri-stimulus time (PST) histogram and rasterplots of the two example neurons, for trials with *B*_*θ*_ = 0.0° (purple) and *B*_*θ*_ = 36.0° (yellow). (**b**) Dynamics of the tuning curves of the same neurons in a 100 ms window starting at the labelled time. (**c**) Scatterplot of the circular variance for the minimum and maximum precision. (**d**) Distributions of the ratio of circular variance between the *B*_*θ*_ = 36.0° and *B*_*θ*_ = 0.0° conditions. (**e**) Distributions of the baseline of the Naka-Rushton functions, *f*_0_, for the two groups of neurons. (ns, not significant; *, *p* < 0.05; ***, *p* < 0.001, Mann-Whitney U-test). (**f**) Distributions of the exponent of the Naka-Rushton functions, *n*. (**g**) Distributions of the spike count distributions of the two groups of neurons, measured as a late (last 100 ms of the trials) over early (first 100 ms of the trials) raw spike count ratio. (**h**) Distributions of the time to maximum tuning curve for the two groups of neurons. (**i**) Average proportion of resilient and vulnerable neurons locations in cortical layers (see Materials and Methods). Error bar represent the standard deviations across animals.

### Recurrent activity controls precision modulations

We then turned to a computational approach to find a mechanistic explanation for the emergence of resilient and vulnerable properties in V1. Mimicking the intra-V1 recurrent connectivity that drives resilient supragranular neurons (12, 18), we modeled the activity of the recorded neurons with a recurrent neural network organized in a “ring” (64) topology (Figure 3a). Such models have been used to explain responses to simple mixtures of oriented inputs as the result of inhibitory activity, which causes non-linear sharpening and decrements of tuning curves, as observed in our data (Figure 1). Indeed, intracortical inhibition plays a central role in shaping the response to stimuli containing multiple orientations, by reducing the activity of the off-median orientation content (3, 11). Electrophysiological experiments have coherently proven that inhibitory processes between oriented components create responses to stimuli of multiple orientation which are smaller than the sum of their parts (25). To account for these considerations, we focused on manipulating only inhibitory activity in our model. It should be noted that inhibition and excitation are balanced variables in cortical networks (17), hence this model is not strictly an argument in favor of an inhibitory-only interpretation of the experimental observations, but rather a useful abstraction about the recurrent nature of the processes involved here.

**Figure 3.**
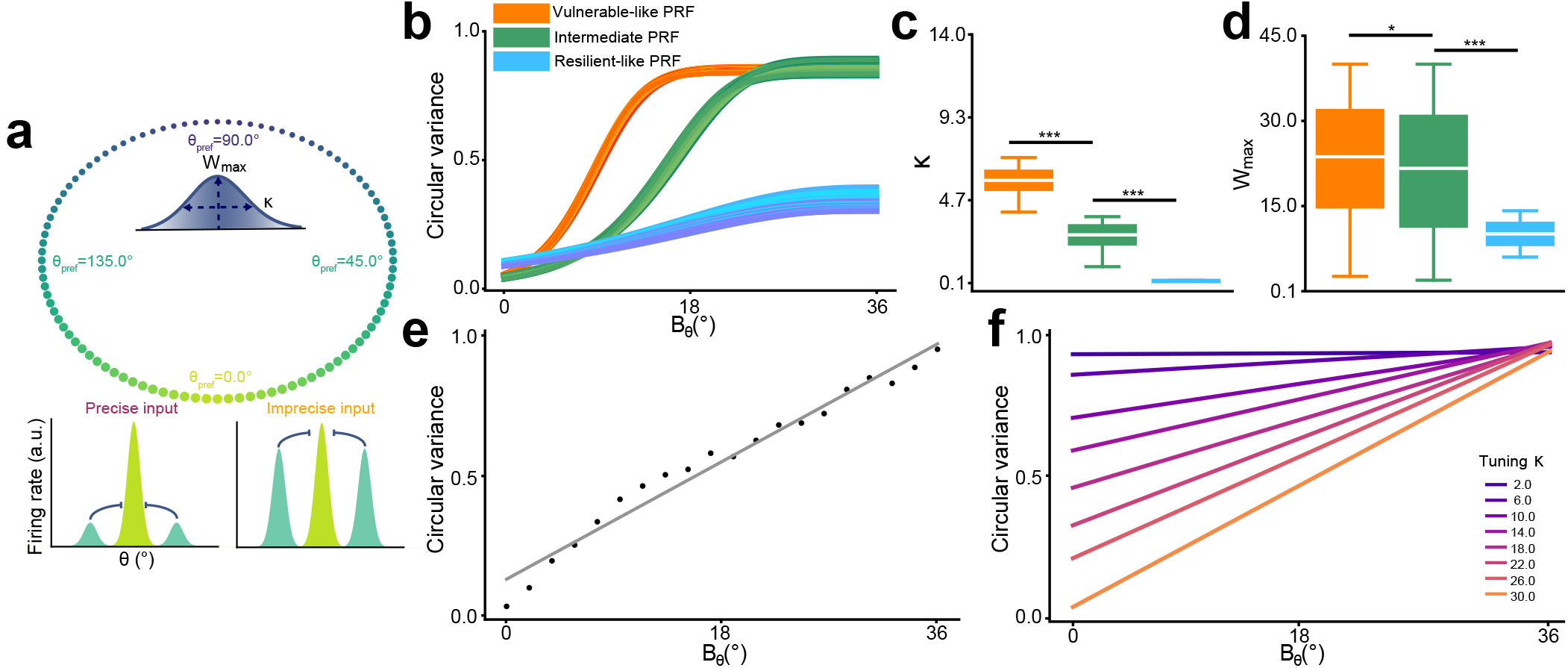
A model of recurrent activity can account for the presence of resilient and vulnerable neurons. (**a**) Top: Illustration of the model. A network of orientation-selective neurons tiling the orientation space is connected by recurrent inhibitory synapses. Two parameters control this connectivity: *W*_max_, the maximum strength of connectivity and *κ*, a measure of the concentration of the activity, with smaller values of *κ* causing a broader extent of the activity in the orientation space (see Materials and Methods). Bottom: Illustration of the effect of precision on V1 activity. Precise inputs (low *B*_*θ*_) elicit an activity centered on few orientation-selective neurons. Conversely, imprecise inputs (high *B*_*θ*_) elicit an activity that spreads to a greater number of neurons, causing an overlap in the recurrent domain of the tuning curves, which drives here an inhibition of the neighboring neurons. (**b**) Selected examples of PRFs with various parameters of simulation. Three different sets of parameters are shown, causing a low *B*_*θ*50_ and high *n* (orange curves), an intermediate *B*_*θ*50_ and high *n* (green curves) and a very high *B*_*θ*50_ and low *n* (blue curves). (**c**) Distribution of the parameter *κ* for these three sets of stimulation. A lower value of *κ* causes a broader inhibition in the orientation space. (***, p < 0.001, Mann-Whitney U-test). (**d**) Distribution of the parameter *W*_max_, the maximum strength of connectivity, for these three sets of stimulations. (*, p < 0.05, Mann-Whitney U-test). (**e**) PRF of the model in a pure feedforward configuration (i.e. *W*_max_ = 0). (**f**) PRF of the model in a pure feedforward configuration, with varying tuning bandwidth of the neurons (‘tuning *κ*’).

Based on the fact that thalamocortical inputs of lower precision increase the spread (in orientation space) of cortical activity (Figure 3a, bottom), we created a phenomenological population rate model of orientation-tuned neurons. In this model, neurons are tiling the orientation space and connect recurrently according to a von Mises distribution (Figure 3a, top), of which we varied two parameters: *W*_max_, the maximum strength of the inhibition and *κ*, a measure of the concentration of the inhibition in orientation space (decreasing *κ* broadens the inhibition, see also Materials and Methods). Varying these model parameters had a non-linear effect on the shape of a neuron’s PRF. By selecting a restricted number of simulations (Figure 3d) which illustrated the prototypical PRFs observed in neuron Ma006 (Figure 1b, top) and Tv001 (Figure 1b, bottom), we found that the transition from the vulnerable to resilient PRF was caused by an increase of the broadness of inhibition in the orientation space (Figure 3c) and a decremented maximum strength of inhibition (Figure 3d).

The range and strength of recurrent activity within a population of orientation-selective neurons is thus sufficient to account for the presence of vulnerable and resilient neurons. Resilient neurons integrate recurrent information over a larger extent of the orientation space, so they should receive a more varied range of orientation-tuned input. While it is known that tuning heterogeneity provides a robust basis to encode mixtures of orientation found in natural images (26), purely feedforward mechanism fail to produce the of nonlinear PRFs observed experimentally (Figure 3e), even with multiple tuning bandwidths in a single population (Figure 3f). In other words, recurrent mechanisms are necessary to reproduce the activity of single neurons. The properties of vulnerable neurons aren’t tied to a specific distance of recurrence, but rather to a specific inhibitory strength which is higher than the one of resilient neurons, fitting the fact that their activity decreases with precision (Figure 1, 2). Based on this modelling approach, it is clear that precision modulations are influenced by multi-neuronal activity, and it thus becomes necessary to understand how V1 populations respond to variations of orientation precision.

### Decoding orientation from population activity

Given that single neuron behavior can be explained by recurrent activity involving multiple neurons, we sought to probe how the global response of the recorded population was modified by variations in input precision. For that purpose, we used a neuronal decoder that probes for population codes in V1, allowing us to understand how orientation and precision were encoded in the overall (recurrent) population activity. The performance of a decoding algorithm is directly tied to the separability of the data in the feature space, that is, the partition of individual tuning curves (6). This is alike to separating multiple oriented components to retrieve the main orientation of the stimulus, which is the process V1 has to accomplish here to perform orientation selectivity. Hence, using a decoding model allows understanding how the neuronal mechanisms studied so far serve V1 in its goal to achieve robust performances when faced with variation of input precision.

We trained a multinomial logistic regression classifier (6), which is a probabilistic model that classifies data belonging to multiple classes (see Materials and Methods). Here, this classifier was fed the firing rate of all recorded neurons in a given time window and learned, for each neuron, a coefficient that best predict the identity (*θ, B*_*θ*_ or *θ* × *B*_*θ*_) of the stimulus (Figure 4a). This classifier was first used to study the changes in the orientation (*θ*) code in the population activity. For this purpose, the dataset of trials was separated for each orientation precision, such that 8 independent, precision-specific, orientation decoders were learned, with optimal parametrization (Supplementary Figure 5). These decoders were then able to predict the correct stimulus orientation, well above the chance level of 1 out of 12 possible orientations. The temporal evolution of decoding performance for these decoders (Figure 4b) showed that maximally accurate orientation encoding is reached faster for stimuli with higher precision (270 and 370 ms post-stimulation for *B*_*θ*_ = 0.0° and *B*_*θ*_ = 36.0°, respectively). It should be noted that as the decoders are trained independently in each time window, the accumulative process observed here is occurring in V1 itself and not in the decoding algorithm. Interestingly, the sustained neural activity present hundreds of milliseconds after the stimulation onset can be used to decode orientation for each precision (Figure 4b,c).

**Figure 4.**
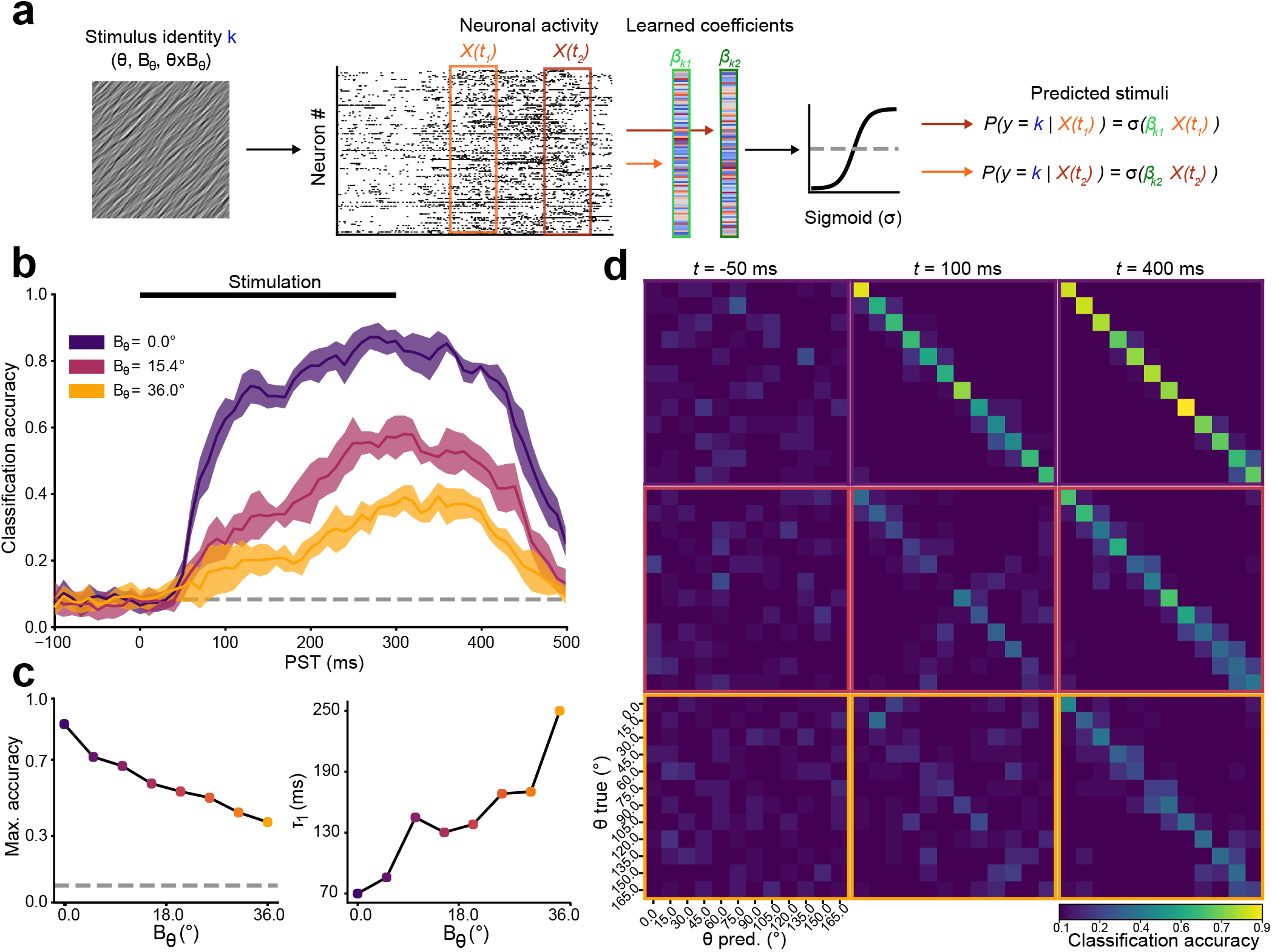
Decoding orientation from population activity. (**a**) Illustration of the decoding model. For each trial, spikes from all neurons in a given time window were pooled in a single neural activity vector *X*(*t*). A multinomial logistic classifier predicts the identity *k* of the stimulus in a given trial by estimating the probability of each hypothesis *y*. This is achieved by learning a coefficient vector *β*_*k*_, which maximizes the accuracy of the prediction 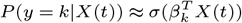, where *σ* is the sigmoid function. The resulting accuracy of this classifier serves as a proxy for the discriminability of the population’s neural activity within feature space. Parameters were controlled (Supplementary Figure 5) to prove that this decoding relied on patterns of spiking activity rather than decoder parametrization. Activity across electrodes and experiments was merged into a single dataset (28, 62), from which the original experiment identity could not be inferred (Supplementary Figure 6). (**b**) Time course around the peri-stimulus time (PST) of the accuracy of three independent decoders trained to predict the orientation *θ* of Motion Clouds for three given orientation precision *B*_*θ*_. Solid dark line represent the mean accuracy of a 5-fold cross validation and filled contour the SD. Decoding at chance level (here,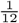) is represented by a gray dashed line. Decoding from resilient and vulnerable neurons (Supplementary Figure 8) showed no difference of maximum accuracy between the two groups. (**c**) Parameters of the decoders’ time courses, estimated by fitting a difference of two sigmoids, with *τ*_1_ the time constant of the first sigmoid (i.e. the time constant of the rising exponential phase of the accuracy curve, see Materials and Methods) (**d**) Confusion matrices of the decoders at three time points around stimulation onset. Values on the diagonals of each matrix represent the instances of correct decoding, that is, where *θ*(*t*)_pred_ = *θ*_true_. Each row of matrices is framed with the color code of the corresponding decoder in (**b**).

Further description of these decoders’ performances is given by confusion matrices (Figure 4d), which represent the accuracy of the decoder for all possible combinations of true and predicted stimuli (see Materials and Methods). The number of significant *y*_*pred*_ = *y*_*true*_ instances for the *B*_*θ*_ = 0° decoder approximately 100 ms after the stimulation onset (upper row) is similar to that of the *B*_*θ*_ = 36.0° decoder (lower row) only reached hundreds of ms later (12/12 versus 11/12 classes significantly decoded above chance level, respectively, *p* < 0.05, 1000 permutations test).

Thus, orientation is encoded in the V1 population code regardless of the oriented input’s precision. The short delay required to process precise inputs is congruent with the feedforward processing latency of V1 (4), while the increased time required to reach maximum accuracy for low precision oriented inputs hints at the presence of a slower mechanism. As was put forward by our model (Figure 3), this could mean a transition from a feedforward-driven regime towards a more recurrent activity as precision of the inputs decrease.

Given the fact that not all neurons are still tuned at low precision (Figure 1), we also examined how resilient and vulnerable neurons contributed to the global population code. Using the same type of decoder, we trained a precision-specific orientation decoder on the spikes of either group of neurons. For precise inputs (Supplementary Figure 8a, left), both groups of neurons reached similar maximum accuracy, which was nonetheless below the combined maximum accuracy reached by decoding from both types of neurons. Unsurprisingly, when precision is low (Supplementary Figure 8a, right), the entirety of the decoding is performed by the activity of resilient neurons, as they are by definition the only neurons still significantly tuned to orientation. These resilient neurons, which have delayed responses (Figure 2), displayed shorter population time constants compared to vulnerable neurons. This could be due to the fact that resilient neurons integrate information from more cortical neurons than vulnerable ones (Figure 3), resulting in each of the spikes carrying more information about the population activity, which increases the performance of the decoder in a shorter time.

### Decoding orientation precision

As the encoding of orientation in V1 was impacted by orientation precision, we next sought to understand if orientation precision itself was encoded in the population activity. A multinomial logistic regression decoder was failed to infer the precision (*B*_*θ*_) of the stimuli from the population activity (Supplementary Figure 7a,b, dark blue). The confusion matrix of this decoder indicates that precise orientations have a distinctive signature (Supplementary Figure 7b, lower row), but once *B*_*θ*_ increases no neuron type can infer the precision of the stimuli (Supplementary Figure 8c). This could stem from the fact that the decoder uses spikes evoked by stimuli of any *θ*. Indeed, as the tuning curves flatten with decrements of precision (Figure 1b, Supplementary Figure 2), it is impossible to distinguish the firing rate at the orthogonal orientation when *B*_*θ*_ = 0.0° from the one at the preferred orientation when *B*_*θ*_ = 36.0°. Thus, encoding orientation precision without any information on the orientation itself is not possible in the current approach. This can be amended by adding an orientation prior to the decoder, by learning only from spikes coming from each neuron’s preferred orientation, which significantly improves precision decoding (Supplementary Figure 7a,b, teal color). Such decoder remained rather inaccurate, partly due to the reduction of the amount of data available for training the algorithm (using only *θ* from the preferred orientation reduced the size of the dataset by 12), which constitutes a machine-learning constraint, rather than a neurobiological limit.

### Decoding orientation and its precision

Thus far, we have shown that precision affects orientation encoding but is not a variable that is explicitly encoded in the population code of V1. We hypothesized that both orientation and its precision could be jointly encoded in V1 as a single variable. We trained a *θ* × *B*_*θ*_ decoder, which could retrieve the identity of Motion Clouds with a maximum accuracy of about 16 times the chance level (1/96) in 350 ms (Figure 5a). This delay to maximum accuracy is similar to the delay observed for low precision decoding of orientation (Figure 4b,c). Unsurprisingly, the confusion matrix of this *θ* × *B*_*θ*_ decoder showed greater accuracy for patterns of higher precision (Figure 5b, 44% average *θ* decoding for *B*_*θ*_ = 0.0° and 12% for *B*_*θ*_ = 36.0°).

**Figure 5.**
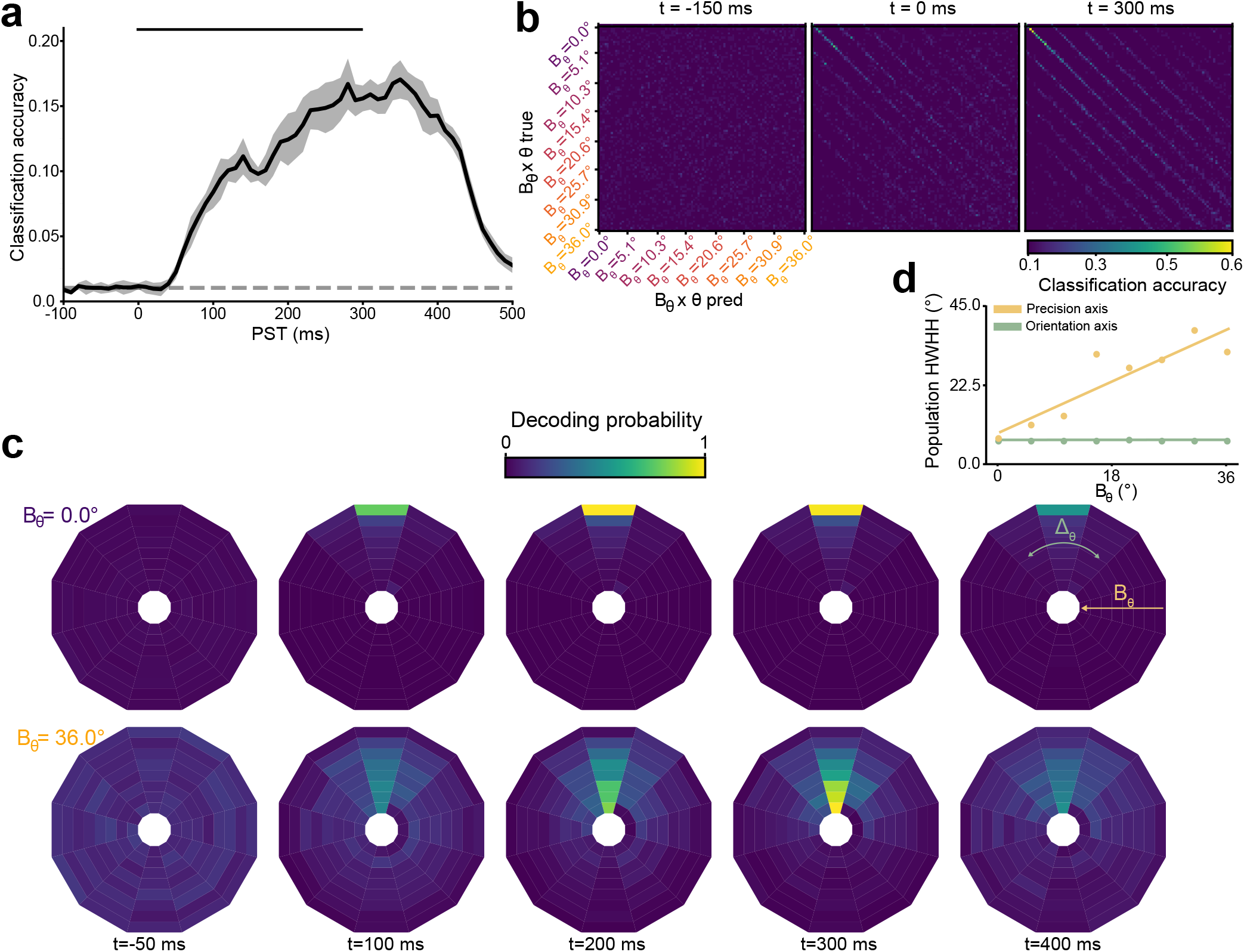
Joint decoding of orientation and precision. (**a**) Time course of a decoder trained to retrieve both the orientation and precision *θ × B*_*θ*_ of Motion Clouds. The solid dark line represents the mean accuracy of a 5-fold cross validation and the filled contour the SD. Decoding at chance level (here, 1/96) is represented by the gray dashed line. (**b**) Confusion matrices of the decoder. In each section labeled by a given *B*_*θ*_, the local matrix organization is identical as in Figure 4, with *θ* = 0.0° located in the upper left corner of the sub-matrix. (**c**) Polar plots of the decoded neural representations. The probabilistic output of the decoder is color-coded in a 2D polar matrix, where the error Δ_*θ*_ on the *θ* identity of the stimulus is represented as the angle of each bin from the upper vertical, while the eccentricity corresponds to the inferred *B*_*θ*_ (lower precision towards the center). The temporal evolution of the probability from the decoder is normalized by row. See Supplementary Figure 9 for a detailed explanation. (**d**) Spreading of population representations. By fitting a von Mises curves on either the orientation (azimuth) or precision (eccentricity) axis of the probabilistic output of the decoder, the Half-Width at Half-Height (HWHH) measures the concentration of the population code around the correct input class. Values of HWHH are fitted with a linear regression, with *y* = .81*x* + 9.85, *r* = .9, p-value = 2.10^−3^ for the precision axis and *y* = 0.0*x* + 7.58, *r* = 0.08, p-value = 0.84 for the orientation axis.

An ambiguity on the inferred precision was observed in these confusion matrices in the form of parallel diagonals (Figure 5b), which are instances where the decoder correctly predicted *θ* but without certainty on the associated *B*_*θ*_ (only elements within the main diagonals are correct decoding occurrences, where *y*_*pred*_ = *y*_*true*_). Rather than assessing the performance of the decoder by reporting the accuracy, which is the argmax of the probability matrix, we lifted this ambiguity on precision by looking at the full probabilistic output of the decoder (see Materials and Methods). In the radially organized representation shown in Figure 5c, the error of the orientation classification, Δ_*θ*_ is represented as the angle of each bin from the upper vertical, while the eccentricity of each bin from the center is representing the precision probability. For instance, in the first row, precisely tuned stimuli (*B*_*θ*_ = 0.0°) elicit right away a probabilistic neural representation of a sharp orientation (low Δ_*θ*_) and precision (no extent on the eccentricity axis) code. By measuring the Half-Width at Half-Height (HWHH) of this probabilistic matrix on either the precision (eccentricity) axis or the orientation (angle) axis, one can observe that as precision decreases (Figure 5c, third row), the spread of the precision code increases, but not the orientation code (Figure 5d). Precision information is thus not absent from the population code, as reported by accuracy metrics (Supplementary Figure 7), but rather encoded in the form of a gradient in the probabilistic representation of the system. Hence, the precision of stimuli is encoded in V1, but in the form of a *θ* × *B*_*θ*_ population code.

Finally, to understand what role does this co-encoding serve, we compared the performances of two orientation decoders: the first learned to decode orientation *θ* without any prior knowledge on precision *B*_*θ*_, while the second was the marginalization of the *θ* × *B*_*θ*_ decoder over *B*_*θ*_, i.e. an orientation decoder with prior knowledge of the precision (Figure 6a). When marginalized, the *θ* × *B*_*θ*_ decoder performed significantly better than the precision-agnostic decoder (12/12 orientation significantly better decoded, *p* < 0.05, 1000 permutations test). Prior knowledge on the orientation precision associated with an oriented input thus yielded better performances of orientation decoding. Training such a marginalized decoder on either group of neurons showed that resilient neurons are significantly better at performing a co-encoding of orientation and precision (Figure 6b). The visualization of the decoder’s coefficients (Figure 6c,d, Supplementary Figure 10) showed how the *θ* × *B*_*θ*_ code is supported by single neurons. These coefficients represent the contributions of each neuron towards the population code: a positive coefficient in a given class *k* means that the neurons’ activity supports the hypothesis of this given class being the correct one, while a negative coefficient would vote against this class. The activity of resilient neurons displayed a pattern coherent with the co-encoding of orientation and its precision, as observed by the extent of the bins in the eccentricity (*B*_*θ*_) axis (Figure 6c,d, bottom). Conversely, the decoding process yielded an orientation-only information from the activity of vulnerable neurons (Figure 6c, top), in which a neuron codes for a single orientation at a single precision.

**Figure 6.**
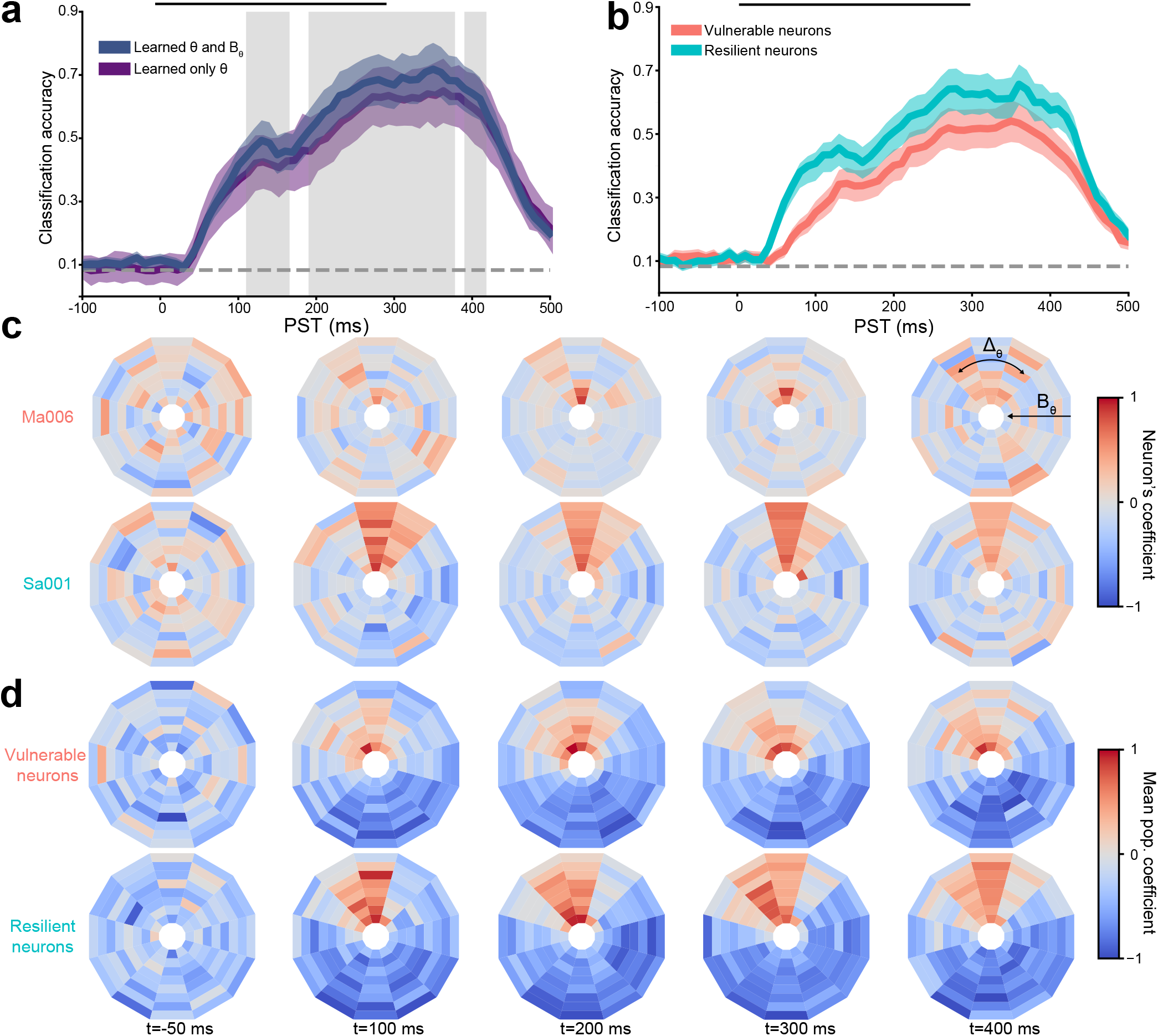
Integrating orientation and precision in a single neural code. (**a**) Time courses of a two independent decoders trained to retrieve orientation *θ*, either by being precision agnostic (red) or by having prior information on precision (blue). The decoder retrieving orientation with prior precision information is a marginalization of the *θ × B*_*θ*_ decoder trained in Figure 5a,b. Solid dark line represent the mean accuracy of a 5-fold cross validation and filled contour the SD. Regions where the marginalized decoder performed significantly better than the precision-agnostic decoder are indicated in grey (1000 permutations test, n=6, threshold p < 0.05). (**b**) Time courses of two marginalized decoders, trained either from vulnerable or resilient neurons’ data. (**c**) Polar plots of the coefficient matrices from a vulnerable (top row) and a resilient (bottom row) neuron. As in Figure 5, the angle of each bin represents the error on the *θ* identity of the stimulus, Δ_*θ*_, and the eccentricity corresponds to the coefficient for each *B*_*θ*_ (lower precision towards the center). Coefficient of a neuron represent a ‘vote’ in the population output, with positive/negative coefficient voting towards/against the associated class *k*, respectively. The temporal evolution of the coefficients from the decoder are normalized for each neuron by row. Additional examples are provided in Supplementary Figure 10. (**d**) Polar plots of the coefficient matrices, averaged across all vulnerable neurons (top row) or all resilient neurons (bottom row).

Taken together, these results can be interpreted in terms of an interplay between cortical populations with dissimilar tuning properties (Figure 7). The accumulative dynamics of the decoders (Figure 4) are directly tied to the notion that neurons are performing incrementally better separation of features within orientation space (6). In other terms, this means that as post-stimulation time elapses, a dominant representation of orientation emerges within the population, which can serve to decode the median orientation contained in the stimuli. Based on the data shown here, we propose that recurrent activity, which mediates precision resilience behaviors (Figure 3), is the biological support of this competition (Figure 7).

**Figure 7.**
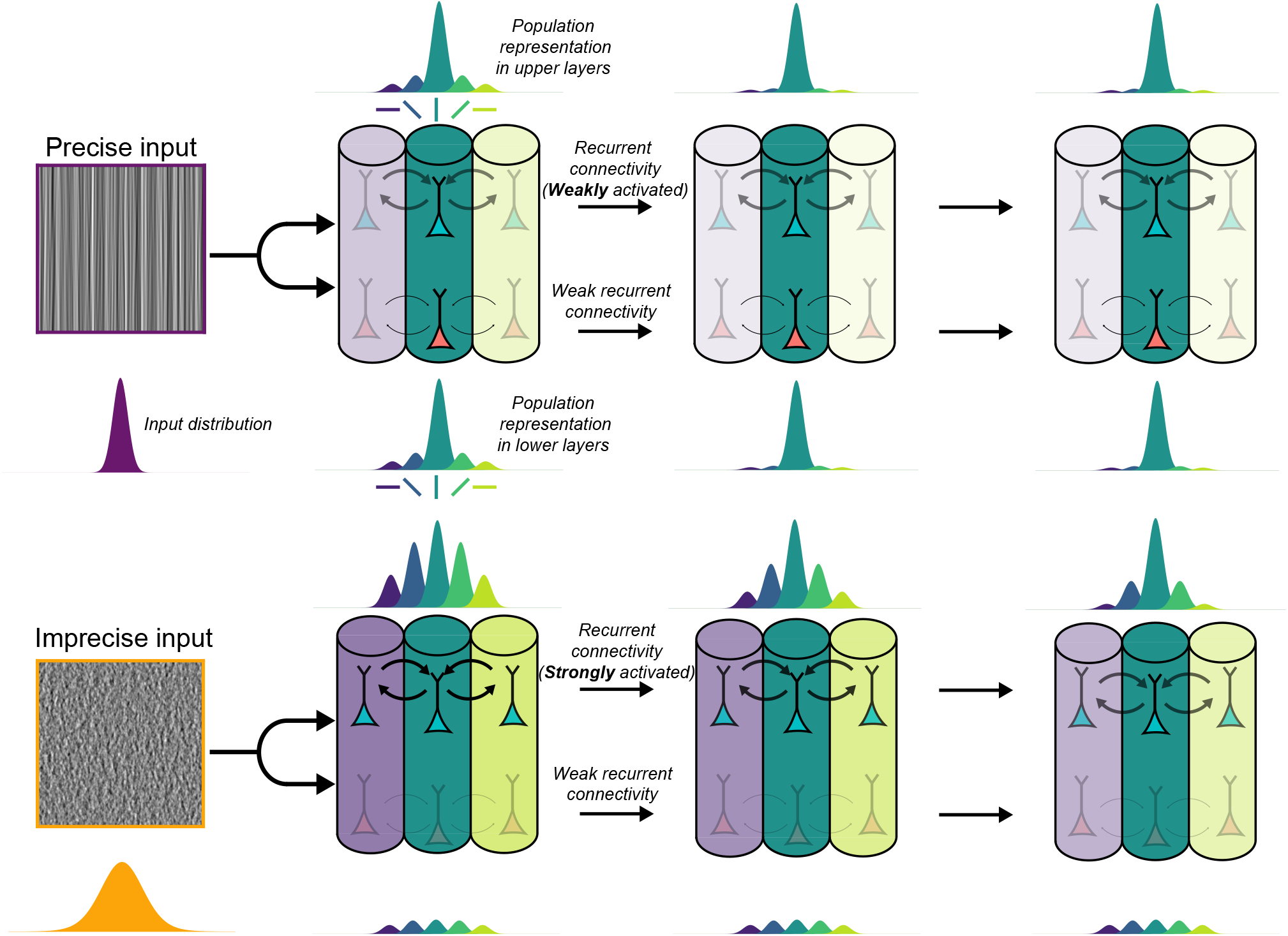
Schematic interpretation of the processing of orientation precision in V1. The color of cortical columns represents their preferred orientation, while the intensity of the color represents the degree of their contribution to the population response. Interlayer connectivity and neuronal types are not shown nor considered here. The underlying distribution of the input as well as the cortical population representations are shown as Gaussian-like curves. Top row: For high precision inputs, the orientation content is highly concentrated around a median orientation, and can be easily read out by V1 with classical feedforward processing, both by resilient (cyan) and vulnerable neurons (orange). Bottom row: For low precision inputs, the distribution is enlarged: retrieving the median orientation requires a competition between multiple components that each needs orientation processing, implicating recurrent connectivity and additional processing time by the supragranular resilient neurons. As processing time goes on, a consensus emerges in the population, which allows maintaining a stable orientation code through multiple level of input precision.

This recurrent process results in a better orientation tuning of resilient neurons, involving slower dynamics (Figure 2). While either group of neuron can encode orientation (Supplementary Figure 8), only resilient neurons can encode precision (Figure 6). The transition from a feedforward-dominated to a recurrent state within the same cortical circuit supports an important property of the population code: as precision of the inputs decreases, the population code remains largely invariant in the orientation domain (Figure 5). This allows V1 to have access to information about orientation and precision, but also creates invariance to variation of precision to support robust orientation encoding at all precision levels.

## Discussion

The precision of oriented inputs impacts orientation selectivity, and we have sought to understand how V1 processes this essential visual feature. Using naturalistic stimuli, we showed that a specific set of neurons responses support a population activity that co-encodes orientation and precision through differential dynamical processes. These findings suggest that recurrent population-level mechanisms in V1 can serve to encode orientation across the varying degrees of precision found in natural images, a process that feedforward mechanisms failed to explain. Here, our main concern was the exploration of feature space, rather than the investigation of spatial relationships. As such, we have used stimuli with no second-order correlation in a full-field setting. Compared to a perfect ecological environment, we have likely recorded sparser cortical responses (29) but also excluded end-stopped cells (24, 35) whose activity could perhaps further help disambiguate imprecise oriented inputs.

We have not pinpointed the neural substrate for precision modulations, but three likely candidates can be put forward, based on theoretical considerations and biological requirements. Precision, by mathematical definition 1*/σ*^2^, requires summing the activity of multiple neurons to compute *σ* at the population level. As shown by the decoding results, this population activity extends over a wide range of time scales, from tens of milliseconds for precise stimuli to few hundreds milliseconds for imprecise ones.

In the classical view of the visual system as a hierarchical network, one first possible substrate of precision processing is the feedback connectivity. In Bayesian formulation, changing the precision of likelihoods (i.e. sensory inputs) drives the posterior closer or further away to the priors (i.e. predictions), with imprecise likelihoods causing the posterior to shift towards the prior (22). One could thus expect that inputs with decreasing levels of precision are encoded by increasingly hierarchically high areas, which would explain the long delays observed for low precision stimuli. Furthermore, even under anesthesia, feedback from extrastriate areas does not change the content of the orientation response in cat V1 (23, 33, 48, 70, 80, 81), but may provide contextual modulation (39), which is arguably similar to precision weighting of cortical activity (41). This explanation seems unlikely due to theoretical considerations, as computing the precision in a cortical area of hierarchical level *N* by another area *N* + 1 creates a non-local process. The brain would face a quickly-scaling problem of regulating cortical flow of information between multiple areas to finely communicate precision without perturbing the communication of cortical information.

One could imagine a simpler, more centralized version of precision processing, where a single brain structure computes the precision of all signals in a given sensory system. In the visual system, such a brain structure could very well be the pulvinar (41), a higher-order thalamic nucleus that establishes cortico-thalamo-cortical connectivity with the visual hierarchy, providing bidirectional shortcuts between visual areas (49). Interestingly, the pulvinar’s action upon the cortex is a function of the hierarchical level of each area (16), regulating and increasing the efficiency of the information transmission across non-linear response functions (14–16). However, computing such precision matrix of any cortical area of size *n* neurons requires an order of *n*^2^ computations, which would be prohibitively costly for the pulvinar (50). It is more likely that the pulvinar could compute an approximation of this precision matrix, which would be then further refined locally within each cortical area. This matches nicely the specific anatomy of transthalamic connectivity, as projections from the pulvinar target supragranular cortical layers (16).

This would lead us to support intra-area cortical recurrence as implementing the specific precision modulation that we have put here in evidence. Predictive coding theories support this interpretation with precision-weighting of predictions errors canonically assigned to supragranular recurrent connectivity (1, 21). In line with our results, recurrent excitatory and inhibitory synaptic connectivity is heterogeneous in V1 (13, 38, 40, 68), which sustains a resilient orientation tuning (51) that could reflect the diversity of single-neurons precision modulations. Temporal scales of recurrent connectivity, namely the slowly-conducted horizontal waves in an orientation map (12), fit the view of precision processing being caused by multiple iterations of a precision computation implemented by recurrent interactions. Furthermore, it would explain the slower population time constant of vulnerable neurons (Supplementary Figure 8a) as the result of a propagation of activity between granular and infragranular layers. While halothane, as used here, increases the latency of visual responses (76), primary sensory cortices are less sensitive to anesthesia than higher-order areas (36, 47). Additionally, intra-V1 recurrent processes observed in anesthetized animals (8, 27) are also frequently reported in awake animals (12, 73). This recurrent interpretation of the results also fits our proposal of low-precision input recruiting more columns of different preferred orientations (Figure 7). Progressive inhibitory suppression of off-median orientation (Figure 4) allows forming a single coherent representation at a given point in the cortical map, rather than an imprecise mixed bag activity.

While there seems to be more arguments towards this last interpretation of our results, we have chosen a conservative approach of our model by considering orientation space rather than actual mapping to neurobiology. This is in part due to the fact that very few other investigations have studied the encoding of orientation precision in V1. A notable exception is from Goris *et al*. (26), which reported that heterogeneously tuned populations are better suited to encode the mixtures of orientations found in natural images. Consistent with their findings, we also reported a variety of single-neuron modulations by the precision of oriented inputs, which then serves to decode orientation mixtures of varying precision at the population level — a requirement for encoding natural images in V1 (55). Their results have further shown that variance of the population activity is directly tied to the (lack of) precision of the stimulation (32). Similarly, we observed that tuning variability (measured as circular variance in Figure 1 and Figure 2) increases for lower precision of sensory inputs. While we do not directly infer precision of stimulus from population variance, we do show that the population variability is a signature of precision, but not of orientation (Figure 5d).

Overall, we have shown that precision, an intrinsic feature of natural images, is actively processed in the primary visual cortex. All three hypotheses about a putative biological substrate create easily testable experimental predictions, which would simply mandate additional electrophysiological recordings in extrastriate areas. This would lead to a better understanding of the neural basis of slow cortical dynamics observed here, which would be a crucial step to properly depart from the fast, feedforward-centric interpretation of visual processes. The use of a generative stimulus framework, as presented here (45) would allow investigating precision modulation at any hierarchical level of the visual system. This rather generic approach could also be extended to other sensory systems, effectively allowing fine control over top-down predictions and bottom-up predictions errors, which could yield pivotal new insights into our understanding of predictive processes in the brain.

## ACKNOWLEDGEMENTS

This article is dedicated to the memory of Umit Keysan (1992-2019), a dear colleague and wonderful friend. The authors would like to thank Genevieve Cyr for her technical assistance, Bruno Oliveira Ferreira de Souza and Visou Ady for advices regarding experimental procedures as well as Louis Eparvier and Jean-Nicolas Jérémie for their comments on the manuscript and Jonathan Vacher for fruitful exchanges on the formalization of the generation of synthetic images and for his contributions to the analysis of neurophysiological recordings. This work was supported by ANR project “Horizontal-V1” ANR-17-CE37-0006 and by a CIHR grant to C.C (PJT-148959). H.J.L. was funded by an École Doctorale 62 PhD grant.

## AUTHOR CONTRIBUTIONS

L.U.P., C.C. F.C., N.C., and H.J.L. designed the study. H.J.L., N.C. and L.K. collected the data. H.J.L., N.C. and L.U.P. analyzed the data. H.J.L. and L.U.P. wrote the original draft of the manuscript. All authors reviewed and edited the manuscript.

## DATA AVAILABILITY

The data generated in the present study are available from the corresponding author, H.J.L., upon reasonable request. No publicly available data was used in this study.

## CODE AVAILABILITY

Data was analyzed using custom Python code, available at https://github.com/hugoladret, using libraries SciPy (79), scikit-learn (58), numpy (78), PyTorch (57), lmfit (54) and Matplotlib (37).

## Materials and Methods

### Experimental Design

Experiments were performed on 3 adult cats (3.6 - 6.0 kg, 2 males). All surgical and experimental procedures were carried out in compliance with the guidelines of the Canadian Council on Animal Care and were approved by the Ethics Committee of the University of Montreal (CDEA #20-006). Animals were first administered atropine (0.1 mg/kg) and acepromazine (Atravet, 1 mg/kg) subcutaneously to reduce the parasympathetic effects of anesthesia and provoke sedation, respectively. Anesthesia was induced with 3.5% Isoflurane in a 50:50 (v/v) mixture of O_2_ and N_2_O. Isoflurane concentration was maintained at 1.5% during surgical procedures. A tracheotomy was performed and animals were immobilized using an intravenous injection of 2% gallamine triethiodide. Animals were then artificially ventilated and a 1:1 (v/v) solution of 2% gallamine triethiodide (10 mg/kg/h) in 5% of dextrose in lactated ringer solution was continuously administered to maintain muscle relaxation. Throughout the experiment, the expired level of CO_2_ was maintained between 35 and 40 mmHg by adjustment of the tidal volume and respiratory rate. Heart rate was monitored, and body temperature was maintained at 37 °C using a feedback-controlled heated blanket. Dexamethasone (4 mg) was administered intramuscularly every 12h to reduce cortical swelling. Pupils were dilated using atropine (Mydriacyl) and nictitaing membranes were retracted using phenylephrine (Midfrin). Rigid contact lenses of appropriate power were used to correct the eyes’ refraction, and eye lubricant was used to avoid corneal dehydratation. Lidocaine hydrochloride (2%) was used in all incisions and pressure points. A craniotomy was performed between Horsley-Clarke coordinates 4-8 P ; 0.5 - 2 L to access the area 17 (V1) contralateral to the stimulation side. Small durectomies were performed for each electrode penetration. A 2% agar solution in saline was applied over the exposed regions to stabilize recordings and avoid the drying of the cortical surface.

### Electrophysiological recordings

During recording sessions, anesthesia was changed to 0.5-1% Halothane, as isoflurane has been shown to yield a depression of visual responses (77). Extracellular activity was recorded using 32 channel linear probes (≈1 MΩ, 1×32-6 mm-100-177, Neuronexus) and acquired at 30 KHz using an Open Ephys acquisition board (71). Single units were isolated using Kilosort (56) and manually curated using the phy software (67). Clusters of low amplitude templates or ill-defined margins were excluded from analysis. Further exclusion was carried if the firing rate of a cluster dropped below 5 spikes.s^−1^ for more than 30 seconds or its tuning curve was poorly fitted with a von Mises distribution (*r*^2^ < .75), leaving 249 putative neurons for further analysis. Clusters were characterized as belonging to supragranular layers if they were located below a fixed distance of 500μm (5) from the top of the electrode (i.e. 5 contacts), which was cross-confirmed with sink/source analysis of the evoked LFP (46).

### Visual Stimulation

Visual stimuli were generated using Psychopy (59) and were projected monocularly with a PROPixx projector (VPixx Technologies Inc., St-Bruno, QC, Canada) onto an isoluminant screen (Da-Lite© screen) located 57 cm from the animal’s eye, covering 104° × 79° of visual angle with a mean luminance of 25 cd/m^2^. The stimuli used here, Motion Clouds (45), are a class of band-pass filtered white-noise textures (10) that provide parametric control over the content of the stimuli while retaining the statistics of natural images (19). The envelope of the filters in Fourier space are Gaussians in the coordinates of the relevant axis, in which they are described by their mean and precision. As such, a Motion Cloud (MC) is defined as:

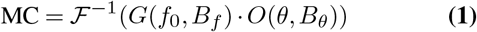

where ℱ is the Fourier transform, *G* the spatial frequency envelope and *O* the orientation envelope. The spatial frequency envelope follows a log-normal distribution:

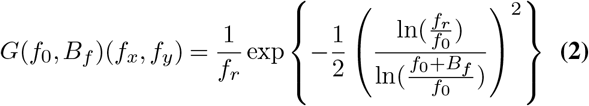

where 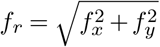 is the radial frequency, f_0_ is the mean spatial frequency and B_f_ is the precision of the spatial frequency distribution, both in cycles per degree. The orientation envelope is a von Mises distribution:

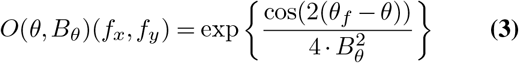

where *θ*_*f*_ is the angle of *f*_*x*_, *f*_*y*_ in the Fourier plane, *θ* is the mean orientation and *B*_*θ*_ is the variance of the orientation distribution. For small *B*_*θ*_, this distribution is close to a Gaussian and *B*_*θ*_ measures its standard deviation. For all stimuli, the spatial frequency parameters were set at *f*_0_ = *B*_*f*_ = 0.9 cpd and a rigid orthogonal drift speed was set to 10°/s, within the response range of area 17 neurons (52). *θ* was varied in 12 even steps from 0 to *π* rad and *B*_*θ*_ in 8 even steps from *π/*5 to ≈ 0. All stimuli were displayed at 100% contrast, for 300 ms, interleaved with the presentation of a mean luminance screen for 150 ms. Trials were fully randomized and each stimulus was presented 15 times.

### Single Neuron Analysis

Tuning curves were constructed by selecting a 300ms window maximizing spike-count variance (74), in which firing rate was averaged and baseline subtracted. Tuning curves were averaged across drift directions and a von Mises distribution (75) was fitted to the data:

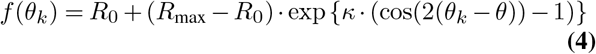

where *R*_max_ is the response at the preferred orientation *θ, R*_0_ the response at the orthogonal orientation, *κ* is a measure of concentration and *θ*_*k*_ the orientation of the stimuli. A global measure of the orientation tuning was assessed by computing the circular variance (CV) of the raw tuning curve data:

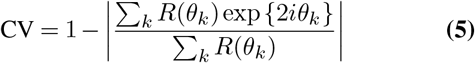

where *R*(*θ*_*k*_) is the response of a neuron to a stimulus of angle *θ*_*k*_. CV varies from 0 for exceptionally orientation-selective neurons to 1 for orientation-untuned neurons (66).

The significance of orientation tuning was assessed by comparing the firing rate of the neuron at the preferred and orthogonal orientations for all trials using a Wilcoxon signedrank test, corrected for continuity. Changes of preferred orientation were assessed as the difference of argmax *f* (*θ*_*k*_) between trials where *B*_*θ*_ = 0.0° and trials where *B*_*θ*_ was great enough so that a neuron was no longer significantly orientation tuned. Variations of peak amplitude were measured by comparing the firing rate at preferred orientation for both these conditions.

### Precision-Response Functions

The variation of circular variance as a function of *B*_*θ*_ was assessed with a NakaRushton function:

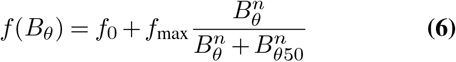

where *f*_0_ is the base value of the function, *f*_max_ its maximal value, *B*_*θ*50_ the orientation precision at half *f*_max_ and *n* a strictly positive exponent of the function (53).

### Computational model

A phenomenological population rate model of a recurrent network was used to reproduce the activity of single neurons. A population of 100 neurons tiling the orientation space in even steps between 0 and *π*rad was modeled as an homogeneous set of orientation-selective units. The orientation distribution of Motion Clouds (Supplementary Figure 1) with 20 precision from *B*_*θ*_ = *π/*5 to ≈ 0 was used to drive the activity of these neurons, according to the equation:

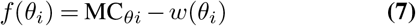

where *f* (*θ*_*i*_) is the activity of a neuron with preferred orientation *θ*_*i*_, MC_*θi*_ the value of the orientation distribution of the input on this orientation and *w*(*θ*_*i*_) the weight of the neighboring inhibitory synapses, described by a von Mises function:

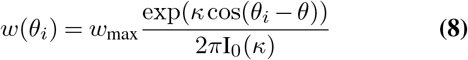

where I_0_(*κ*) the Bessel function of order 0 and, *θ*_*i*_ is a neighboring neuron with index *i, w*_max_ the maximum synaptic weight and *κ* the measure of concentration of the function. The effect of *w*_max_ and *κ* were explored by varying both parameters with 60 values, between 0.0; 40.0 and 0.0; 15.0, respectively. Furthermore, a non-linear threshold was added to the response of each neuron, such that neuron who fell beneath this threshold failed to produce a spiking activity. This parameter was not varied, and an empirical value of ≈ 10% of the maximum input to the model (when *B*_*θ*_ ≈ 0) was used.

### Population Decoding

The stimulus categorical identity was decoded using a multinomial logistic regression classifier (6), which can be interpreted as an instantaneous, neurally plausible, readout of the population activity (4). For a given stimulus, this population activity was a vector *X*(*t*) = [*X*_1_(*t*) *X*_2_(*t*) … *X*_249_(*t*)], where *X*_*i*_(*t*) is the spike count of neuron *i* in a time window [*t*; *t* + Δ*T* ], where *t* is the time bin and Δ*T* the size of the integration window, usually 100 ms. Time *t* was varied from −200 ms to 400 ms, relative to the stimulation onset, in steps of 10 ms. Each time bin was labeled as the end of the time window and was hence reported as *t* +Δ*T*.

Practically, the multinomial logistic regression is an extension of the binary logistic regression (6), which was here trained to classify the neural activity vector between *K* possible classes. The probability of any such vector to belong to a given class is:

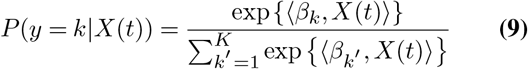

where ⟨·,·⟩ is the scalar product over the different neurons, *k* = 1, …, *K* is the class out of *K* possible values and *β*_*k*_ are the vectors of learned coefficients of the classifier. We trained several such classifiers, to decode orientation *θ* (*K* = 12), orientation precision *B*_*θ*_ (*K* = 8) or both (*K* = 12 × 8 = 96). All meta-parameters (the integration window size, penalty type, regularization strength and train/test split size) were controlled, showing that the decoder is optimally parameterized and that its performances result from experimental data and not decoder parametrization (Supplementary Figure 5). Decoding accuracy was reported as the average accuracy across classes, also known as the balanced accuracy score (9), which accounts for possible imbalances in the learning or testing datasets. Decoding performance was also reported in the form of confusion matrices, in which the values on each row *i* and column *j* represents the normalized number of times a stimulus of class *k* = *i* is predicted to belong to the class *k* = *j*. Hence, a perfect decoder would produce a perfectly diagonal confusion matrix (unit matrix). We reported the significance of decoders by splitting the population activity in separate training and testing data sets, then performing 6 different such splits on different groups of neurons, comparing the resulting confusion matrices to class-shuffled decoder matrices. Previous publications have reported population decoding by merging neural activity across electrodes or experiments (28, 62), which we validated in our data (Supplementary Figure 6).

### Decoding time course

To estimate the temporal parameters of the decoders, the evolution of the population decoding accuracy through time was fitted with a difference of two sigmoid functions:

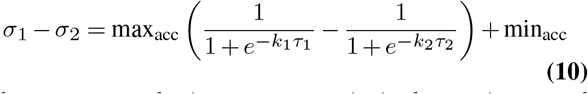

where max_acc_ and min_acc_ are respectively the maximum and minimum accuracies of the decoder, *k* the steepness and *τ* the time constant of the function.

### Statistical Analysis

All data were prepared for analysis with custom Python code. Statistical analysis was performed using non-parametric tests. Wilcoxon signed-rank test with discarding of zero-differences was used for paired samples and Mann-Whitney U test with exact computation of the *U* distribution was used for independent samples. Due to the impracticality of using error bars when plotting time series, colored contours are used to represent SEM values, with a solid line representing mean values. In boxplots, the box extends from the lower to upper quartile values, with a white line at median. The upper and lower whiskers extend to respectively *Q*1 − 1.5 * *IQR* and *Q*3 + 1.5 * *IQR*, where *Q*1 and *Q*3 are the lower and upper quartiles and *IQR* is the inter-quartile r

## Figures

**Supplementary Figure 1.**
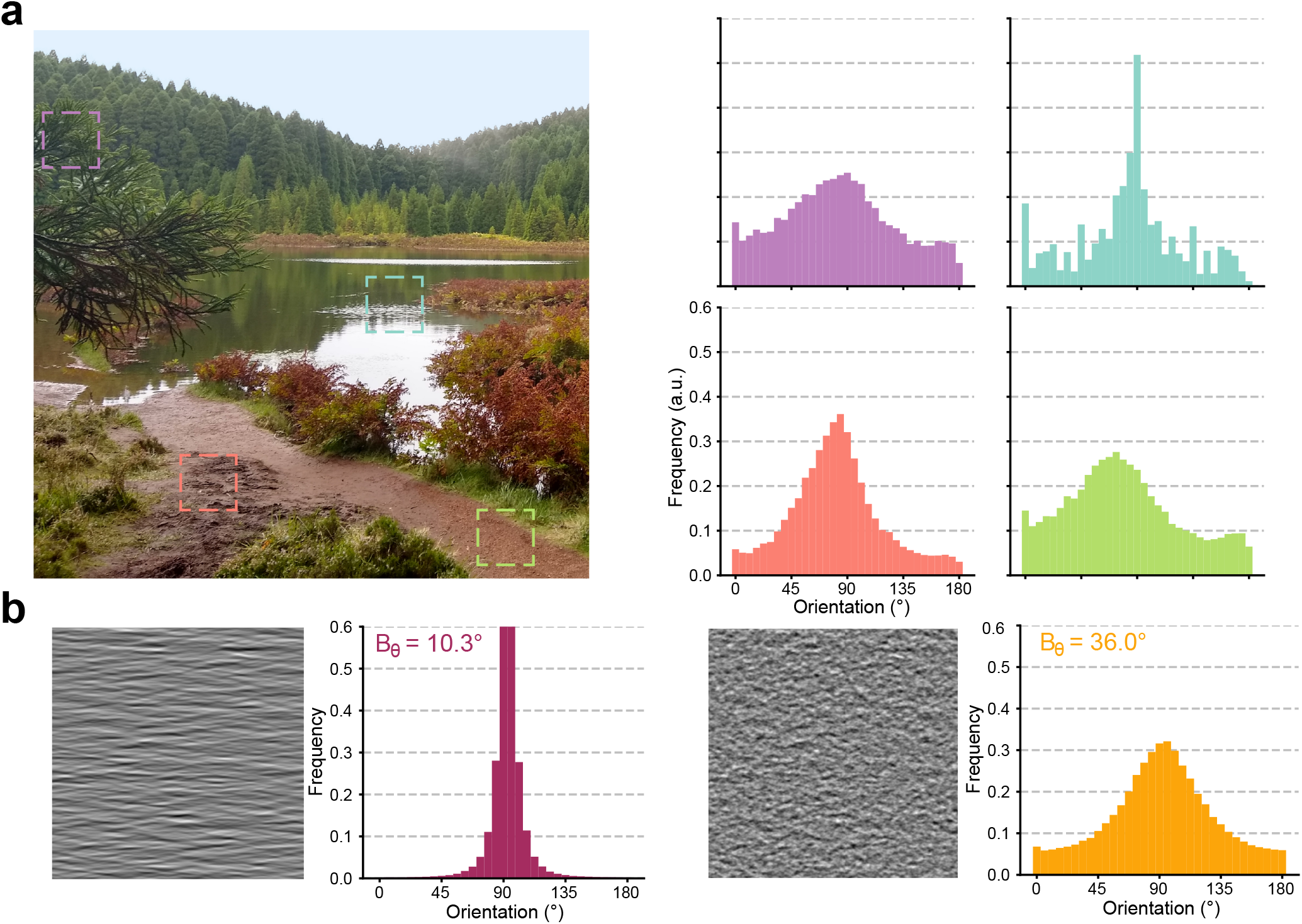
Orientation composition of natural images and natural-like stimuli. (**a**) Distributions of orientations retrieved from a natural image (São Miguel Island, Azores, Portugal, picture taken by H.J.L.) using an Histogram of Oriented Gradients (16×16 pixels cell size, 200×200 pixels image subsets, color-coded to the associated distributions). (**b**) Single frames from Motion Clouds stimulus, oriented at an angle of 90° relative to the vertical axis, of high (*B*_*θ*_ = 10.3°) and lowest precision (*B*_*θ*_ = 36.0°).

**Supplementary Figure 2.**
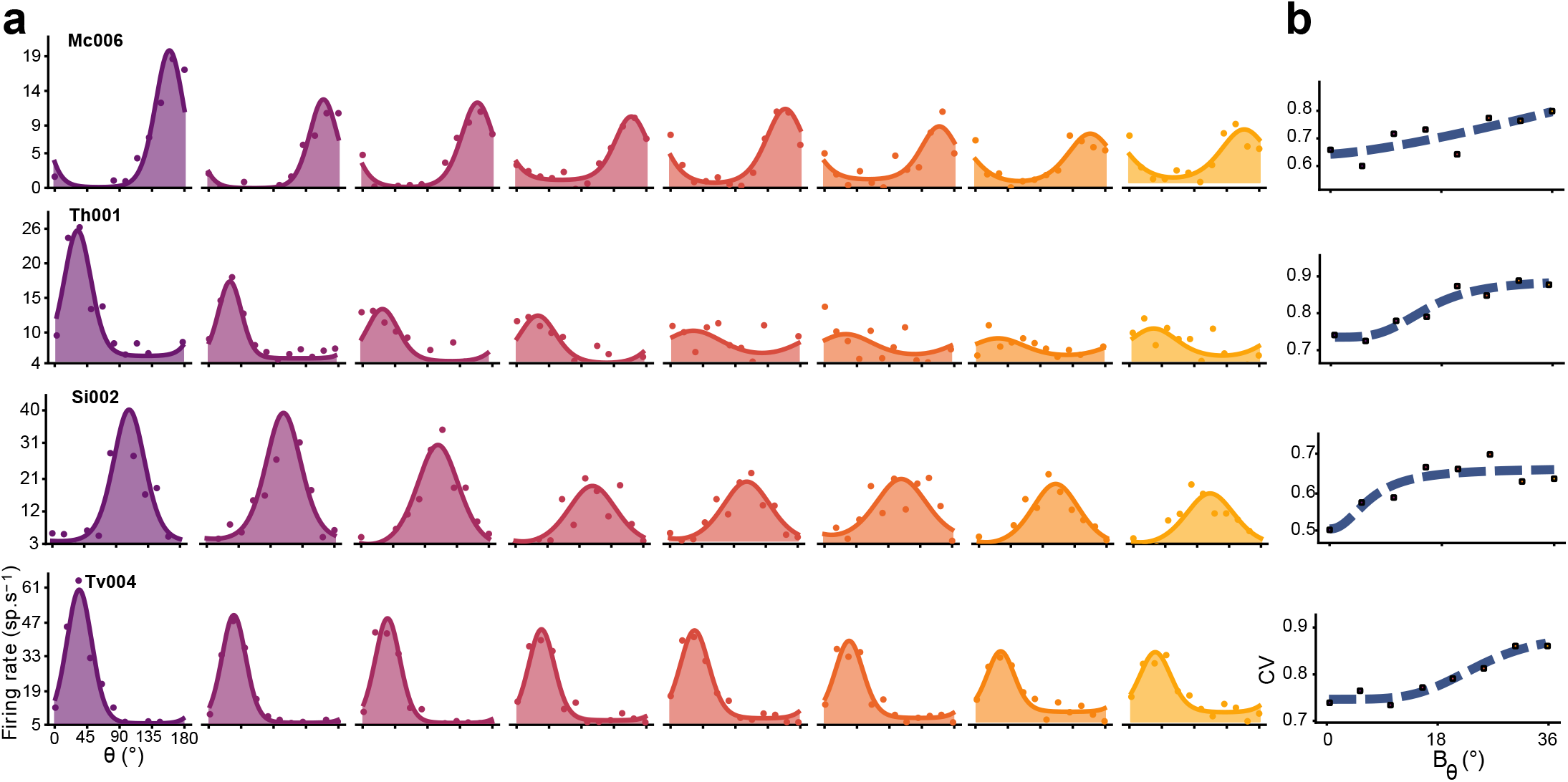
Additional examples of single neuron tuning curves and PRFs. (**a**) Tuning curves of 4 additional neurons in response to Motion Clouds of increasing *B*_*θ*_ (left to right). Colored points represent the mean firing rate across trials (baseline subtracted, 300 ms average), with lines indicating a fitted von Mises function. (**b**) Precision-response functions (PRFs), assessing the changes of orientation tuning as measured by the Circular Variance (CV) as a function of *B*_*θ*_, fitted with Naka-Rushton (NKR) functions (dashed curves).

**Supplementary Figure 3.**
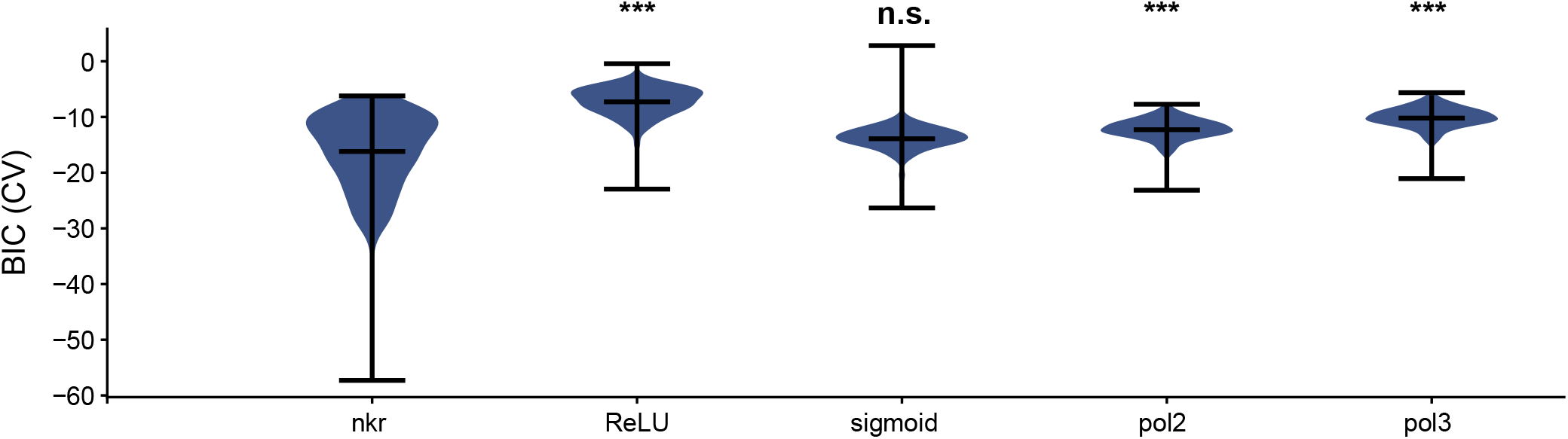
Criterion on the model selection for fitting the PRFs. Violin plot of the Bayesian Information Criterion (BIC) of the CV curves of all recorded neurons. Each violin plot represent a different type of fitted equation, respectively: Naka-Rushton (`nkr`, see Materials and Methods) ; Rectified Linear Unit (`ReLU`, *f* (*x*) = *max*(0, *x*)) ; logistic function 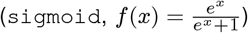 ; second degree polynomial function (`pol2`, *f* (*x*) = *ax*^2^ + *bx* + *c*) and third degree polynomial function (`pol3`, *f* (*x*) = *ax*^3^ + *bx*^2^ + *cx* + *d*). A lower BIC indicates less information lost in the fitting process, hence a better fitting model. Naka-Rushton curves were chosen over sigmoid functions for the skewness of the BIC distributions towards negative values, as well as the explainability of their parameters. `n.s.`, not significant; *, p < 0.05; **, p < 0.01; ***, p < 0.001 (Kruskal-Wallis H-test, post-hoc Dunn Pairwise test, Bonferroni corrected).

**Supplementary Figure 4.**
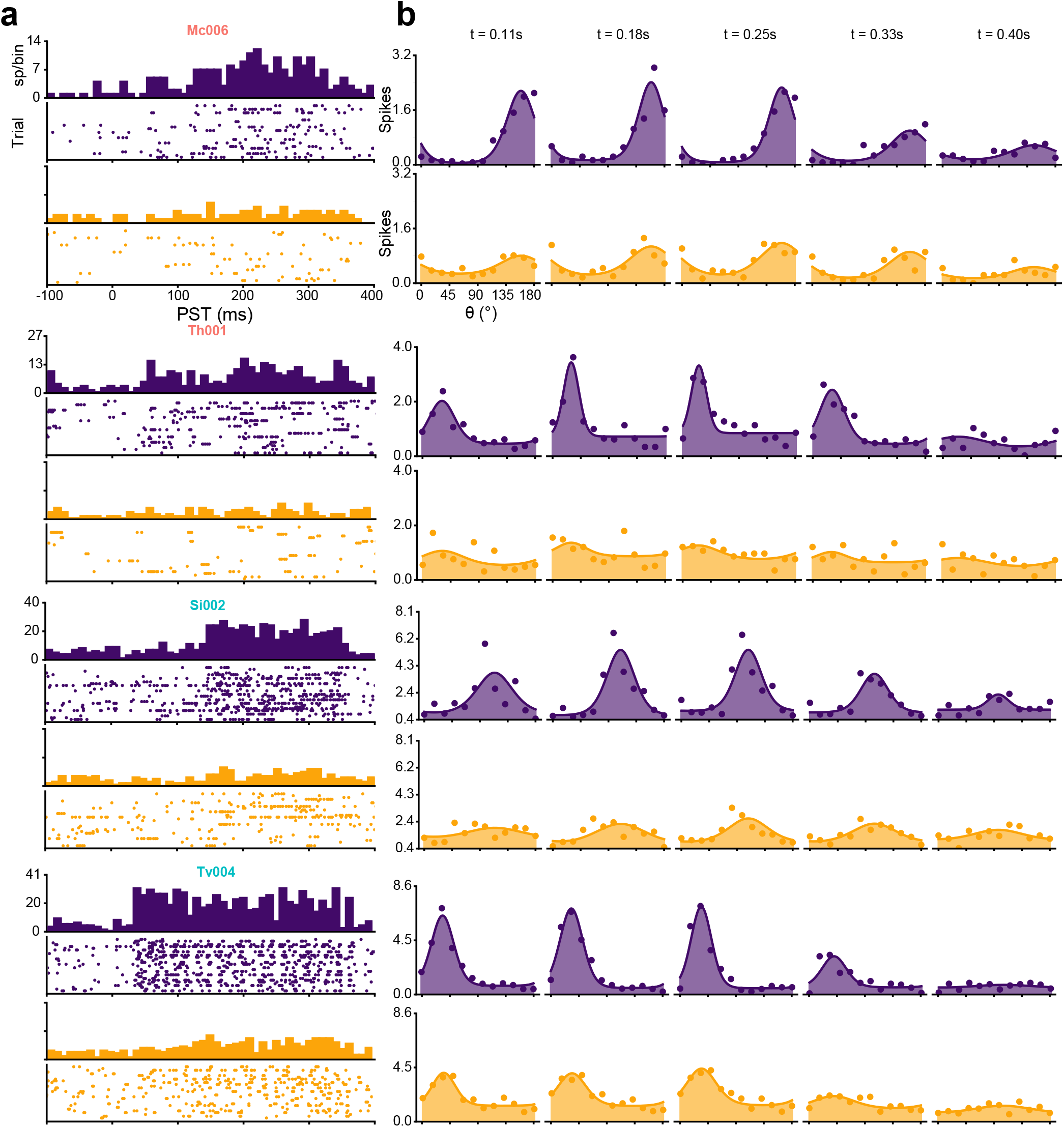
Additional examples of dynamical responses of neurons. Neuron labels are colored in red for neurons untuned to stimuli with *B*_*θ*_ = 36.0° (“vulnerable neurons”), and conversely in teal when still significantly tuned to the same low-precision stimuli (“resilient neurons”) (**a**) Peri-stimulus time (PST) histogram and rasterplots of the example neurons, for trials with *B*_*θ*_ = 0.0° (purple) and *B*_*θ*_ = 36.0° (yellow). (**b**) Temporal evolution of the tuning curves of the same neurons, taken in a 100 ms window after the reported time.

**Supplementary Figure 5.**
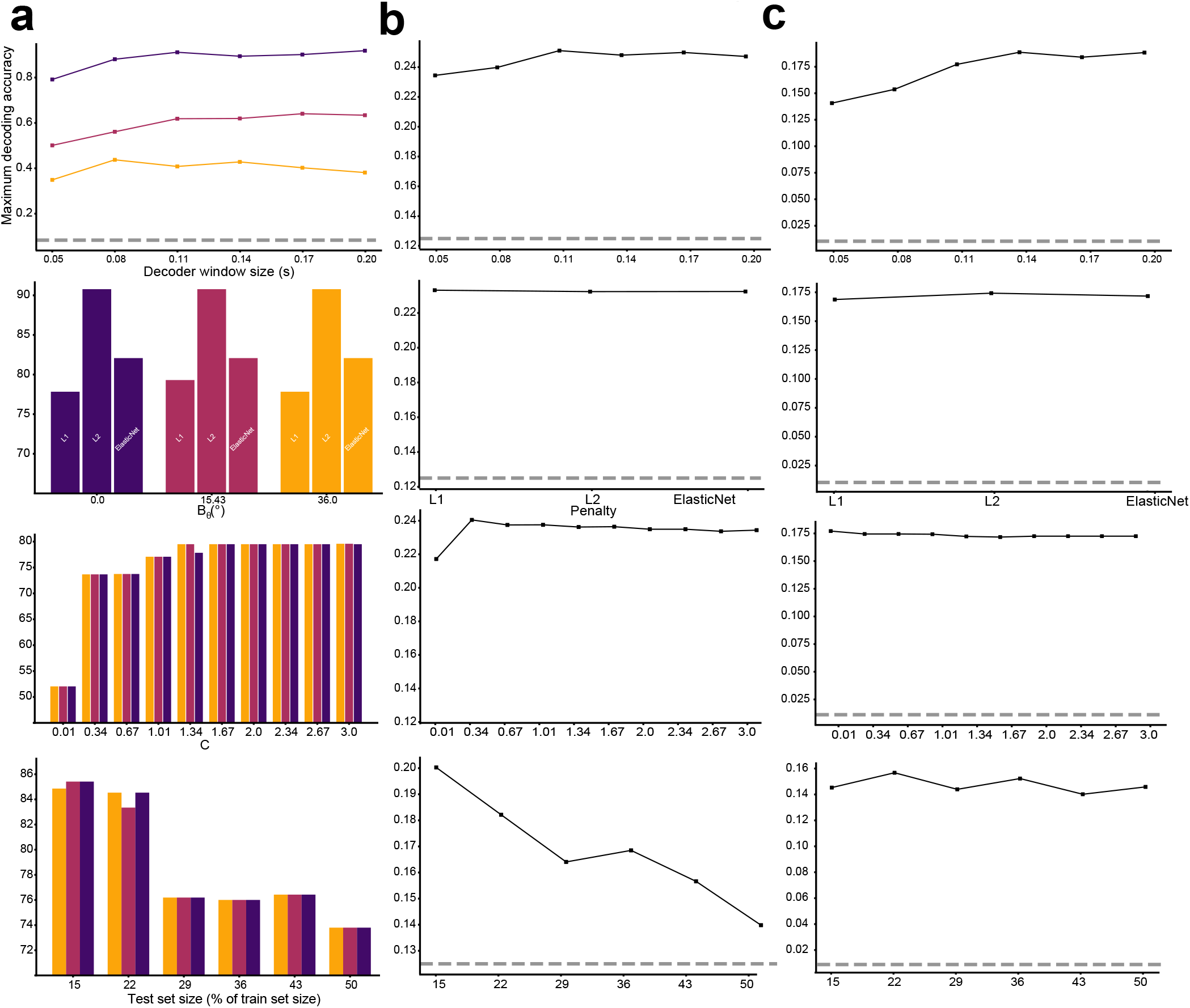
Parameters evaluation of the decoders. (**a**) Parameter scanning of the orientation *θ* decoders for three *B*_*θ*_. Maximum accuracy reached during the time course of each decoder was reported for each parameter. From top to bottom, the parameters optimized are: the length of the time window Δ*T*, the type of penalization norm applied to the decoder, the regularization strength parameter and the percentage of data kept out of the training set to evaluate the decoder’s accuracy. (**b**) Parameters of the orientation precision *B*_*θ*_ decoder. (**c**) Parameters of the orientation and orientation precision *B*_*θ*_ *× θ* decoder.

**Supplementary Figure 6.**
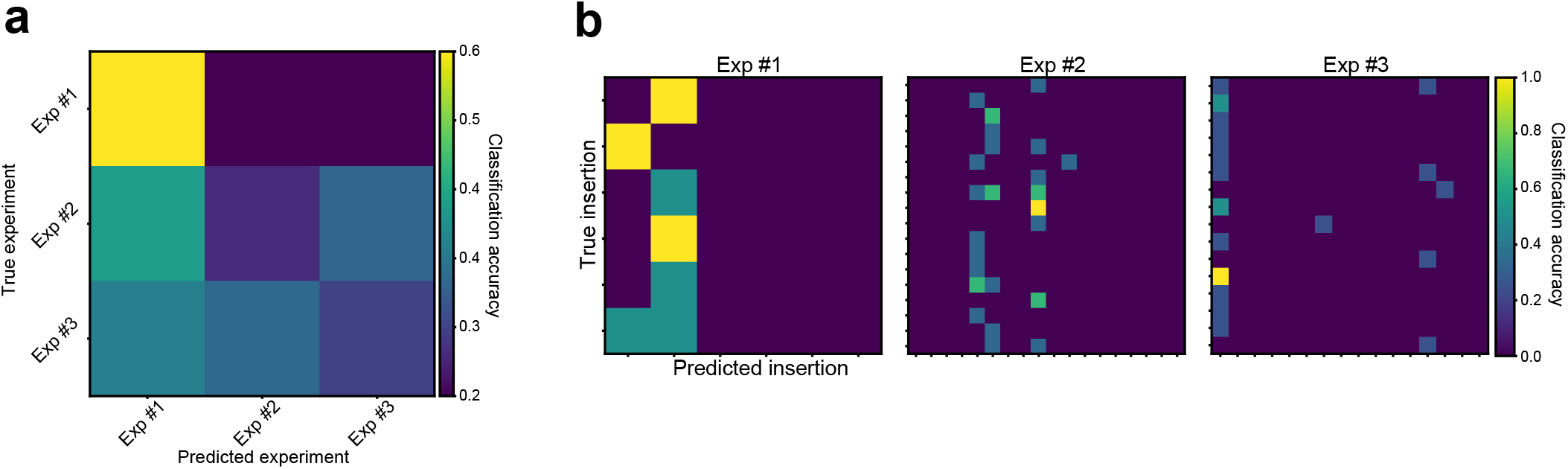
Decoding the experiment identity and insertion identity of neurons. (**a**) Confusion matrix of a decoder trained to retrieve the experiment identity using three groups of 30 neurons (bootstrapped 1000 times). (**b**) Confusion matrices of three decoders trained to retrieve the insertion identity of the neurons recorded in each experiments.

**Supplementary Figure 7.**
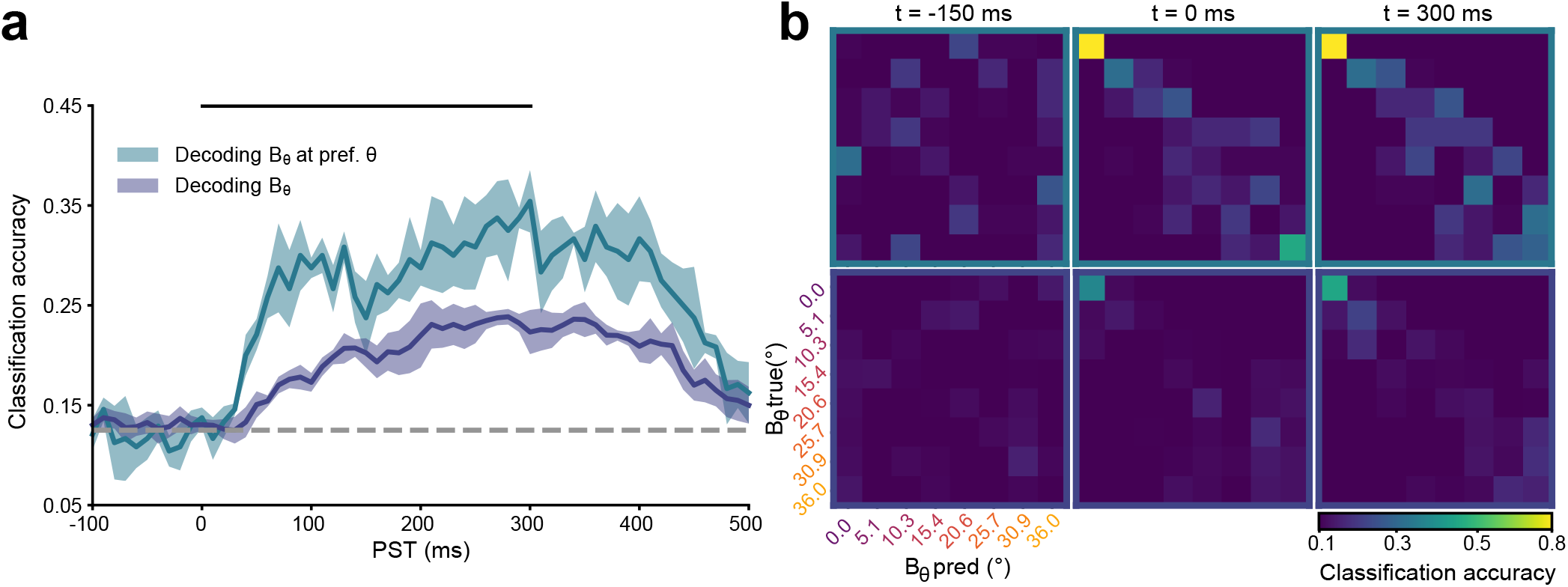
Decoding orientation precision from population activity. (**a**) Time course of two independent decoders, trained to retrieve the orientation precision *B*_*θ*_ of Motion Clouds either with (teal) or without (dark blue) orientation *θ* prior. Solid dark line represent the mean accuracy of a 5-fold cross validation and filled contour the SD. Decoding at chance level (here, 1/8) is represented by a gray dashed line. (**b**) Confusion matrices of the two decoders, for the decoding task with orientation prior (top) and without (bottom).

**Supplementary Figure 8.**
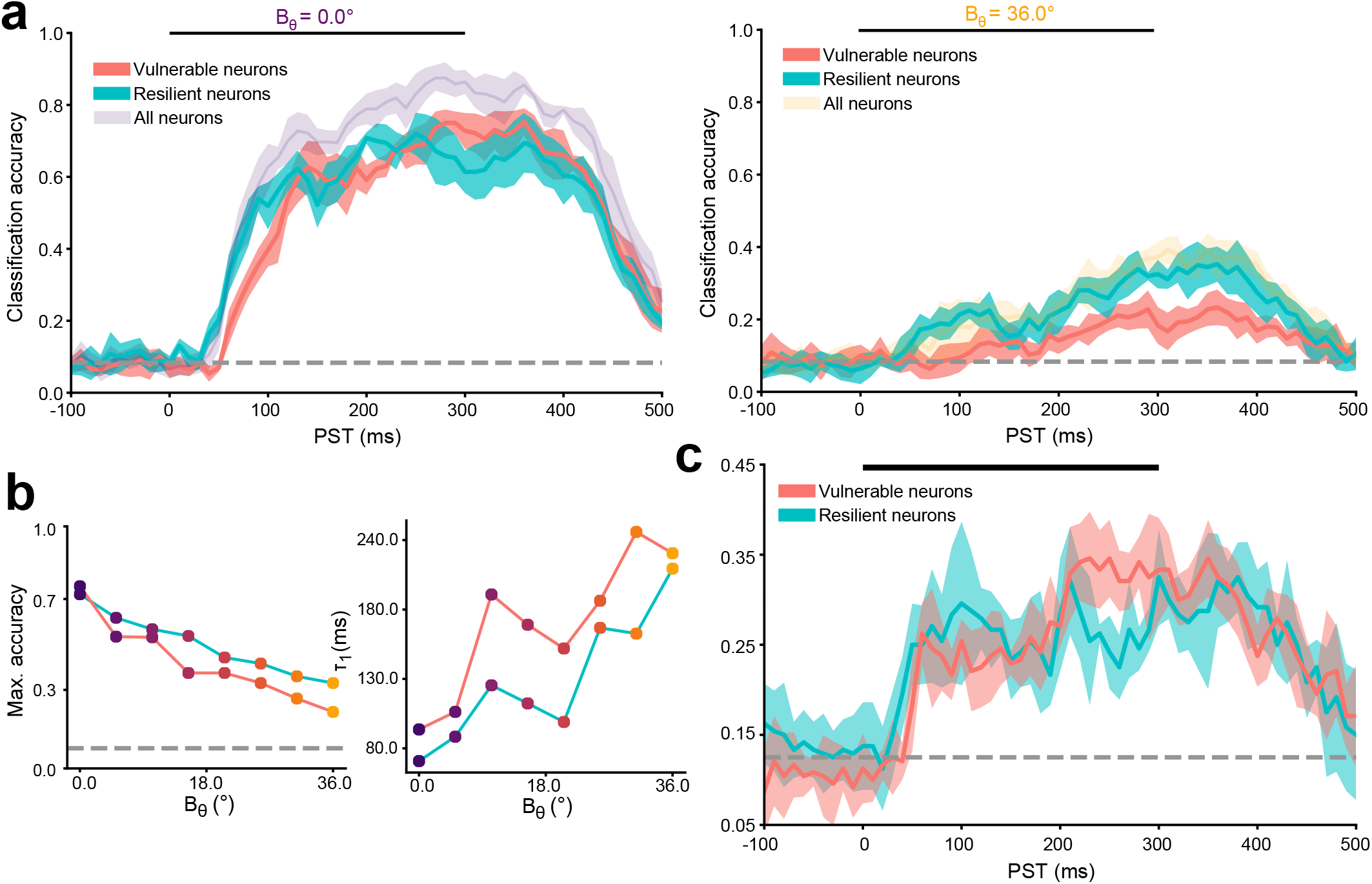
Decoding stimulation properties from vulnerable and resilient neurons. (**a**) Left: Time course around the peri-stimulus time (PST) of the accuracy of three independent decoders, trained to predict the orientation *θ* of maximally precise Motion Clouds from either vulnerable, resilient, or all neurons (identical to Figure 4b). Right: Time course of three independent decoders, trained on the same task but for Motion Clouds of the lowest precision. Solid dark line represent the mean accuracy of a 5-fold cross validation and filled contour the SD. Decoding at chance level (here, 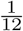) is represented by a gray dashed line. (**b**) Parameters of the decoders’ time courses trained from vulnerable and resilient neurons’ data, estimated by fitting a difference of two sigmoids. (**c**) Time course around the PST of the accuracy of two decoders, trained to retrieve the precision of the stimulation with the preferred *θ* prior (identically to Supplementary Figure 7b,teal), from either vulnerable or resilient neurons.

**Supplementary Figure 9.**
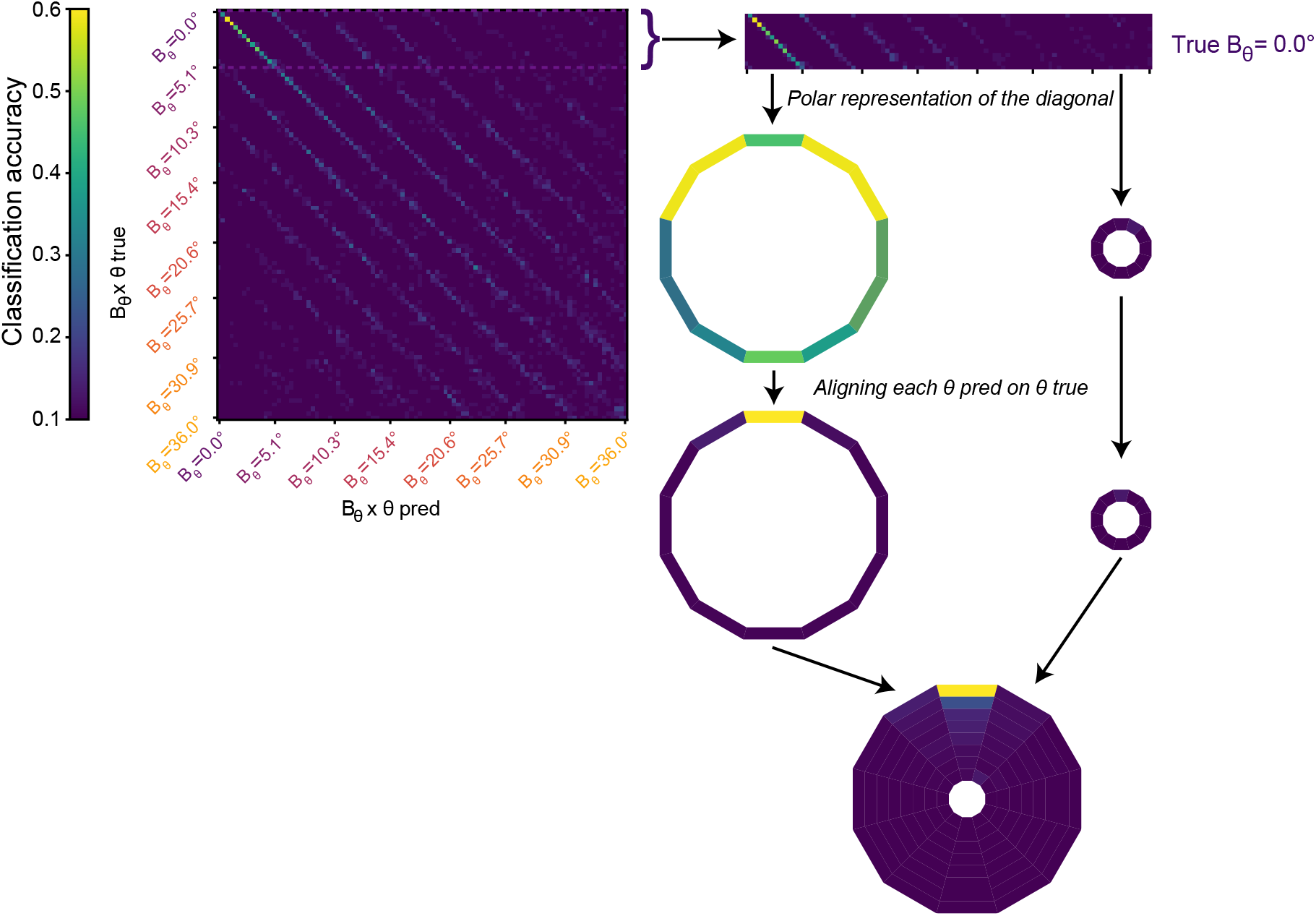
Schematic representation of the process used to obtain polar representations from the *θ × B*_*θ*_ decoder’s confusion matrices. For each row, in which all stimuli decoded belong to one given precision *B*_*θ*_, the diagonal from each possible column, in which all stimuli are inferred as belonging to one of the 8 possible *B*_*θ*_ is taken and aligned over the true *θ*. This process is repeated for each row, which produces a full polar representation of the decoding output. When representing coefficient matrices, as in Figure 6, the initial data comes from coefficient vectors *β*_*k*_ (see materials and methods).

**Supplementary Figure 10.**
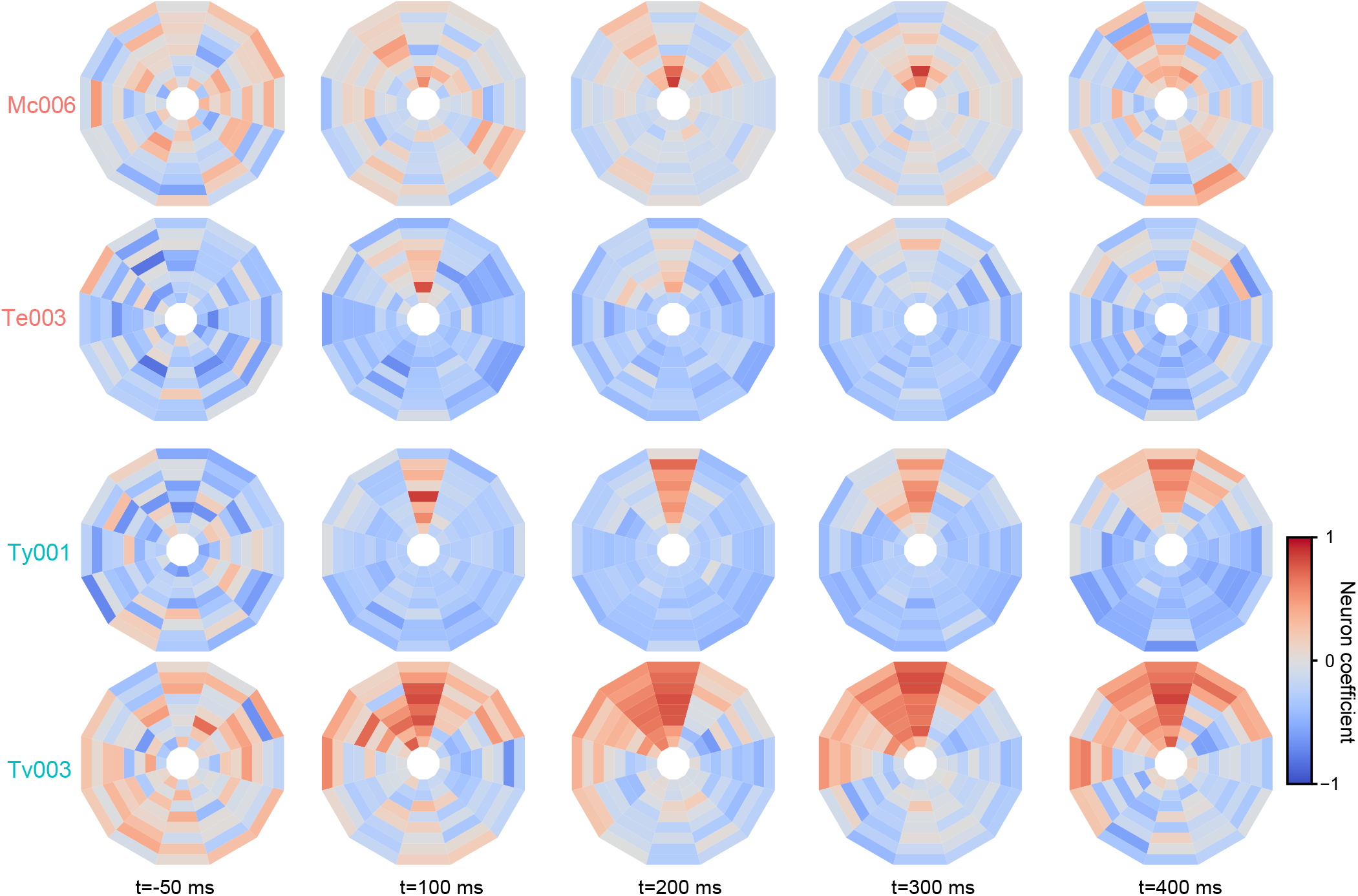
Polar plot of coefficient matrices from additional neurons from two vulnerable (top rows) and two resilient (bottom rows) neurons. As in Figure 5, the angle of each bin represents the error on the *θ* identity of the stimulus, Δ_*θ*_, and the eccentricity corresponds to the coefficient for each *B*_*θ*_ (lower precision towards the center). The temporal evolution of the coefficients from the decoder are normalized for each neuron by row.

